# Lineage Tracing Reveals a Shared Cellular Origin for Supraclavicular Brown and Inguinal Beige Adipocytes

**DOI:** 10.1101/2025.10.02.680118

**Authors:** Yali Ran, Kai Zhang, Yi-Ting Shen, Qianxing Mo, Ziyi Wang, Hari Krishna Yalamanchili, Sharon John, Mari Kogiso, Xia Gao, Chunmei Wang, Tanvi Sinha, Brian L Black, Miao-Hsueh Chen

**Author notes:** Corresponding Author Miao-Hsueh Chen, Ph.D. Children’s Nutrition Research Center, Department of Pediatrics, Baylor College of Medicine 1100 Bates, St. rm 10018 Houston, TX, 77030, USA.

## Abstract

The metabolic importance of brown adipose tissue (BAT) has been recognized, but the origins of BAT, particularly supraclavicular BAT (scBAT), are less clear. Here, we traced the origin of scBAT to Mef2c-anterior heart field (AHF)-marked cells. Mef2c-AHF-marked cells isolated from scBAT can spontaneously differentiate into brown adipocytes, express mesenchymal stem cell markers, and can be isolated from the stromal-vascular fraction (SVF) of wild-type scBAT as [CD31^-^CD45^-^SCA-1^+^CD29^+^CD34^-^CD24^-^] (CD34^-^) cells. Mef2c-AHF-marked cells substantially overlap with Prrx1-marked cells in scBAT, which also contribute to beige adipocytes in inguinal white adipose tissue (iWAT) during development. Similarly, CD34^-^ cells isolated from the SVF of iWAT can spontaneously differentiate into beige adipocytes *in vitro*. Using intersectional tracing, we revealed that common progenitors of Mef2c-AHF- and Prrx1-marked cells in scBAT emerge from the developing heart. Thus, these studies reveal a common origin for scBAT and cardiac tissue and a close lineage relationship between scBAT and beige adipocytes of the iWAT.

## Introduction

Brown adipose tissue (BAT) is crucial for metabolism. BAT consists of brown adipocytes that store small lipid droplets, have high mitochondrial densities, and generate heat through non-shivering thermogenesis^1,2^. Recently, beige adipocytes, similar in morphology and thermogenic potential to brown adipocytes, were found in inguinal white adipose tissue (iWAT)^3,4^.

Because of its unique function of converting fat and other nutrients into heat, BAT is considered a potential therapeutic candidate for treating cardiometabolic diseases, such as obesity and type II diabetes^5,6^. Recent studies show that BAT exists not only in small animals and infants, but also in adult humans^7–10^. Unlike most mouse BAT depots, including the interscapular, subscapular, and auxiliary BAT depots located in the dorsal thoracic region, the largest human BAT depot, supraclavicular BAT (scBAT), is located above the clavicle bone. To understand human BAT function, we and others identified a previously unknown BAT depot located in the mouse neck that anatomically resembles human scBAT^11–13^. Morphologically, mouse scBAT consists of loosely connected thin strips of brown adipocyte islands^12^. Molecularly, supraclavicular brown adipocytes express high levels of *Ucp1* and have high thermogenic potential^12,13^.

Brown adipocytes arise from brown adipocyte progenitor cells earlier than white adipocytes during embryonic development. Genetic lineage tracing studies revealed that brown adipocytes in the dorsal thoracic region, specifically the interscapular BAT (iBAT), arise from Myf5-marked progenitor cells^14^. *Myf5*, a transcription factor, is expressed in the paraxial mesoderm that gives rise to the myotomal compartment of the somites, which will eventually give rise to skeletal muscles of the body^15,16^. Additional genes that also mark the interscapular brown adipocyte progenitor cells include *En1*, *Pax3*, and to a lesser extent *Pax7*, which are all expressed in somite during early developmental stages^17–20^. Based on these tracing studies, brown adipocytes and myoblasts are thought to descend from common somite-derived cells^20^. While genetic tracing is not applicable in humans, studies using next generation sequencing have identified an array of markers for human brown adipocytes^3,21–26^. Interestingly, a significant portion of these markers are also expressed in mouse beige adipocytes, suggesting human brown adipocytes are molecularly similar to the mouse beige adipocytes^3,24,25^. Thus, a consensus on whether human brown adipocytes are classic brown adipocytes or newly identified beige adipocytes has not been definitively reached.

Since the rediscovery of adult human brown adipocytes over a decade ago, the function and importance of human brown adipocytes have begun to be revealed^5,27–31^. However, at least two critical questions remain unresolved. First, the origin of scBAT is unknown. Second, the lineage relationship between supraclavicular brown and beige adipocytes has not been investigated.

To address these unresolved questions in the present study, we applied the Cre-loxP-mediated lineage tracing method to determine that supraclavicular brown adipocytes originate from Mef2c-AHF-marked (Mef2c-AHF^+^) cells in scBAT. We also show that Mef2c-AHF traced-cells differentially express several mesenchymal stem cell markers. Using these markers, we establish a panel of cell surface markers for isolating Mef2c-AHF^+^ supraclavicular brown adipocyte progenitors from scBAT in wild-type mice. In addition to being marked by the Mef2c-AHF-Cre driver, supraclavicular brown adipocytes were also marked by the Prrx1-Cre driver, which is known to mark adipocytes, including beige adipocytes, in iWAT. Applying the same panel of cell surface markers for isolating Mef2c-AHF^+^ cells, we isolated beige adipocytes from iWAT from wild-type mice. These studies indicate a close lineage relationship between scBAT and iWAT, including beige adipocytes. Lastly, intersectional tracing of both Mef2c-AHF- and Prrx1-marked cells revealed that Mef2c-AHF- and Prrx1-marked cells largely overlap in the primordium of scBAT, and these cells appeared to emerge from the developing heart.

## Results

### scBAT does not originate from somite-derived cells

In mice, a thin layer of scBAT is situated between the lower jaw and the clavicle bone, spreading symmetrically from the ventral medial (ventral scBAT) toward the lateral side of the neck (lateral scBAT) (Figure 1A). To probe the origin of the scBAT, we first tested whether this depot arises from somites, like iBAT does. We used the Myf5^Cre^ driver, where Cre is expressed in somite-derived cells during development, including BAT depots in the dorsal thoracic trunk^15^. We bred *Myf5^Cre^* mice with *R26R-LacZ* (R26R) mice to produce *Myf5^Cre^;R26R* embryos (Figure S1A) and performed X-gal staining to visualize LacZ-positive cells. X-gal-stained cells were visible in the somite of embryonic day 10.5 (E10.5) *Myf5^Cre^;R26R* embryos (Figure S1B). At E18.5, when muscles and BAT depots had formed, only iBAT stained positive for β-galactosidase (Figure S1C). To extend our tracing to postnatal mice, we used three Cre drivers: *Myf5^Cre^*, *Pax7^Cre^*, and *Pax3^Cre^*. Like *Myf5^Cre^*, *Pax7^Cre^* and *Pax3^Cre^*mice mark somite-derived cells, including BAT depots^18,19^. These Cre mice were bred with dual fluorescent protein *ROSA^mTmG^* reporter mice to produce the following progeny: *Myf5^Cre^;ROSA^mTmG^*, *Pax7^Cre^;ROSA^mTmG^*, and *Pax3^Cre^;ROSA^mTmG^* (Figure 1B). To visualize Cre-marked membrane-targeted GFP-positive (mG^+^) brown adipocytes, iBAT, scBAT, iWAT, and epididymal WAT (eWAT) were isolated from these mice at 3 to 5 weeks of age. As previously reported, nearly all the brown adipocytes were mG^+^ in iBAT isolated from *Myf5^Cre^;ROSA^mTmG^* and *Pax3^Cre^;ROSA^mTmG^*mice, and to a lesser extent from *Pax7^Cre^;ROSA^mTmG^*mice (Figure 1C). Scattered mG^+^ cells were also detected in iWAT from *Myf5^Cre^;ROSA^mTmG^*and *Pax3^Cre^;ROSA^mTmG^* mice (Figure 1C). In contrast, mG^+^ brown adipocytes were absent from scBAT from *Myf5^Cre^;ROSA^mTmG^* and *Pax7^Cre^;ROSA^mTmG^* mice, and only a few mG^+^ brown adipocytes were seen in scBAT from the *Pax3^Cre^;ROSA^mTmG^*mice (Figure. 1C). These findings indicated that, unlike iBAT, scBAT does not originate from somite-derived cells.

**Figure 1.**
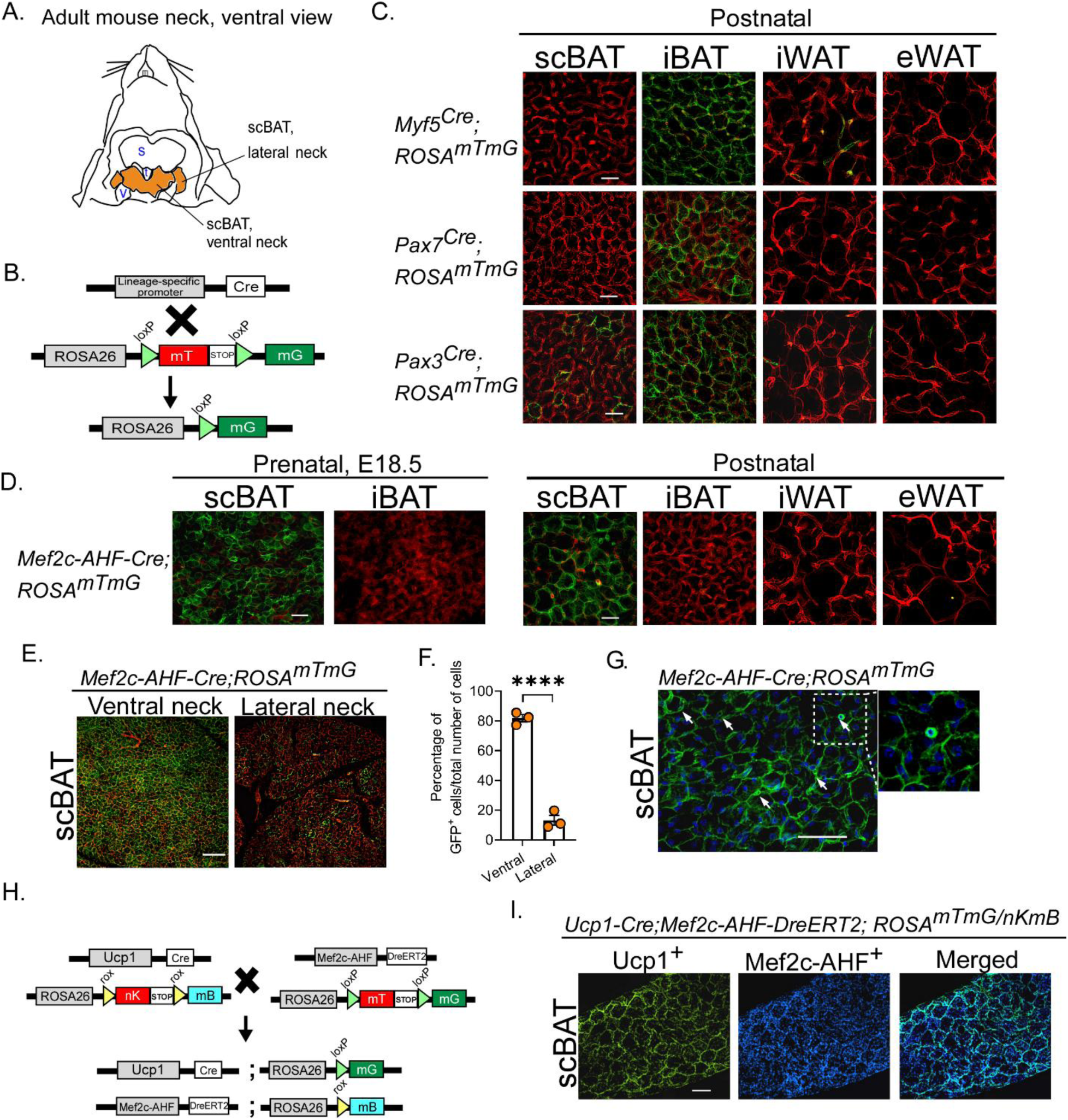
Mef2c-AHF-Cre driver marks cells in scBAT. (A) A diagram describing the relative anatomical location of mouse scBAT. s: salivary gland. t: trachea. v: jugular vein. (B) A schematic diagram of the lineage tracing strategy using *Myf5^Cre^*, *Pax7^Cre^*, and *Pax3^Cre^* mice crossed with *ROSA^mTmG^* mice to identify somite-derived brown adipocytes. (C) Representative sections of scBAT, iBAT, iWAT, and eWAT from *Myf5^Cre^;ROSA^mTmG^*, *Pax7^Cre^;ROSA^mTmG^*, and *Pax3^Cre^;ROSA^mTmG^* mice. Scale bar = 25 μm. n = 2-4. (D) Representative sections of scBAT, iBAT, iWAT, and eWAT in prenatal (E18.5) and postnatal 3-week-old *Mef2c-AHF-Cre;ROSA^mTmG^* mice. Scale bar = 25 μm. n = 2-4. (E) Representative images of Mef2c-AHF-marked cells in scBAT from the ventral and lateral neck regions of postnatal 3-5-week-old *Mef2c-AHF-Cre;ROSA^mTmG^*mice. Scale bar = 100 μm. n = 3. (F) The percentage of mG^+^ cells from ventral and lateral neck region of scBAT. Data are presented as mean ± SEM. ****, P < 0.0001. n = 3. (G) A higher resolution image of Mef2c-AHF-marked cells in the scBAT of a 3-week-old *Mef2c-AHF- Cre;ROSA^mTmG^* pup. Arrows point to small round shaped mG^+^ cells. Scale bar = 50 μm. n = 3. (H) A schematic diagram of the double lineage tracing strategy used to generate *Ucp1-Cre*;*Mef2c- AHF-DreERT2;ROSA^mTmG/nKmB^* mice. (I) Representative images of the expression of Ucp1- and Mef2c-AHF-marked cells in scBAT of the postnatal *Ucp1-Cre*;*Mef2c-AHF- DreERT2;ROSA^mTmG^/^nKmB^* mice. Scale bar = 25. μm. n = 2.

### Supraclavicular brown adipocytes originate from cardiac progenitor-derived cells

The neck, where both mouse and human scBAT are found, is a complex anatomical region connecting the body and head. The progenitor cells contributing to neck anatomical structures can emerge from the primordium of the head or heart. Recent studies have shown that some neck muscles and blood vessels share a common origin with the heart^32,33^. Given that scBAT is positioned near the sternocleidomastoid muscle and jugular veins in the ventral neck, we hypothesized that scBAT could originate from cells that also contribute to the developing heart. To test this, we used *Mef2c-anterior heart field (AHF)-Cre* mice, where Cre expression is driven by an enhancer fragment from the Mef2c transcription factor-encoding gene-in AHF derivatives^34^. We bred *Mef2c-AHF-Cre* mice with *R26R* mice to produce *Mef2c-AHF-Cre;R26R* embryos and performed X-gal staining. We confirmed that Cre only marked the developing heart in E10.5 *Mef2c-AHF-Cre;R26R* embryos (Figure S1D). At E18.5, only scBAT was positive for β-galactosidase, but not iBAT, providing the first evidence that scBAT originated from Mef2c-AHF-marked cells (Figure S1E). To confirm this observation, we generated *Mef2c-AHF-Cre;ROSA^mTmG^* mice. We sectioned E18.5 *Mef2c-AHF-Cre;ROSA^mTmG^* fetuses and scBAT, iBAT, iWAT, and eWAT from *Mef2c-AHF-Cre;ROSA^mTmG^* mice at 3-week-old to visualize Mef2c-AHF-marked cells. Again, mG^+^ cells were detected in scBAT and absent from other depots, further demonstrating that scBAT but not iBAT descends from Mef2c-AHF-marked cells (Figure 1D).

Interestingly, mG^+^ cells were not uniformly distributed in scBAT. Rather, these cells were highly expressed in the ventral scBAT and to a much lesser extent in the lateral scBAT (Figures 1E&F). We also noticed that large and small round shaped mG^+^ cells were each present in scBAT (Figure 1G), suggesting that Cre could be expressed in both mature brown adipocytes and progenitor cells. To verify that the Cre marked *Ucp1* expressing adipocytes, we employed a dual-recombinase lineage (Cre-loxP/Dre-rox) tracing system, utilizing the *Mef2c-AHF-DreERT2* and *Ucp1-Cre* driver mice to mark Mef2c-AHF- and Ucp1-dual-positive cells (Figure 1H). In this tracing system, Dre can be activated by injecting Tamoxifen (Tam) at the time that *Ucp1* is highly expressed in postnatal mice. The Mef2c-AHF Dre-marked cells are revealed by *ROSA^nKmB^* mice carrying Dre recombination target sites, the rox sequences^35^. We generated *Ucp1-Cre;Mef2c-AHF-DreERT2;ROSA^nKmB/mTmG^* mice and activated Dre by injecting Tam 5 weeks after birth. Cre-marked Ucp1-positive adipocytes overlapped with Mef2c-AHF-marked cells in scBAT (Figure 1I), confirming the presence of Mef2c-AHF- and Ucp1-dual-positive brown adipocytes in scBAT.

### Mef2c-AHF^+^ cells can spontaneously differentiate into mature brown adipocytes *in vitro*

To directly test whether Mef2c-AHF-marked cells give rise to brown adipocytes in scBAT, we dissociated the SVF from the scBAT of *Mef2c-AHF-Cre;ROSA^mTmG^* mice at age 2.5-to 5-week-old and applied fluorescence-activated cell sorting (FACS) to isolate mG^+^ cells from the SVF (Figures 2A&B). These sorted mG^+^ cells are referred to as Mef2c-AHF^+^ cells hereafter to reflect their origin. Freshly isolated Mef2c-AHF^+^ cells were cultured and expanded once or twice prior to administering adipogenic differentiation assays using standard adipogenic induction medium (IM) or a combination of Bmp7 and Rosiglitazone (Rosi), two master regulators of brown adipogenesis (Figure 2C). Surprisingly, during expansion of these sorted cells, we noticed that the shape of some sorted Mef2c-AHF^+^ cells gradually changed from triangular or rectangular to round adipocytes containing small lipid droplets (Figure 2D). Oil-Red-O and LipidSpot 610 staining of neutral lipids confirmed the appearance of lipid droplets in Mef2c-AHF^+^ cells (Figure 2D), indicating these Mef2c-AHF^+^ cells are likely stem/progenitor cells capable of spontaneous differentiation *in vitro*. Induction with IM or a combination of Bmp7 (conditioned medium) and Rosi (Bmp7+Rosi) induced full differentiation of Mef2c-AHF^+^ cells in just 8 days after induction as indicated by abundant Oil-Red-O-and LipidSpot 610-stained neutral lipids (Figure 2E). To confirm that these differentiated Mef2c-AHF^+^ cells were brown adipocytes, we performed RT-qPCR to measure the expression levels of genes involved in brown adipogenesis, including *PPARg*, *Fabp4*, *Pgc-1α*, *Ucp1*, and *β3-AR* (beta3 adrenergic receptor), and two mitochondrial marker genes, *Nrf-1*, and *Tfam,* in Mef2c-AHF^+^ cells that differentiated spontaneously or were induced by IM or by Bmp7+Rosi. As shown in Figure 2F, all these genes were expressed in the Mef2c-AHF^+^ cells. The expression levels of *Pgc-1a* and *Ucp1*, two thermogenic marker genes, significantly increased after stimulation with Isoproterenol (Iso), a beta-adrenergic receptor agonist. Together, these *in vitro* analyses indicated both spontaneously and adipogenic induced Mef2c-AHF^+^ cells gave rise to mature brown adipocytes with thermogenic potential.

**Figure 2.**
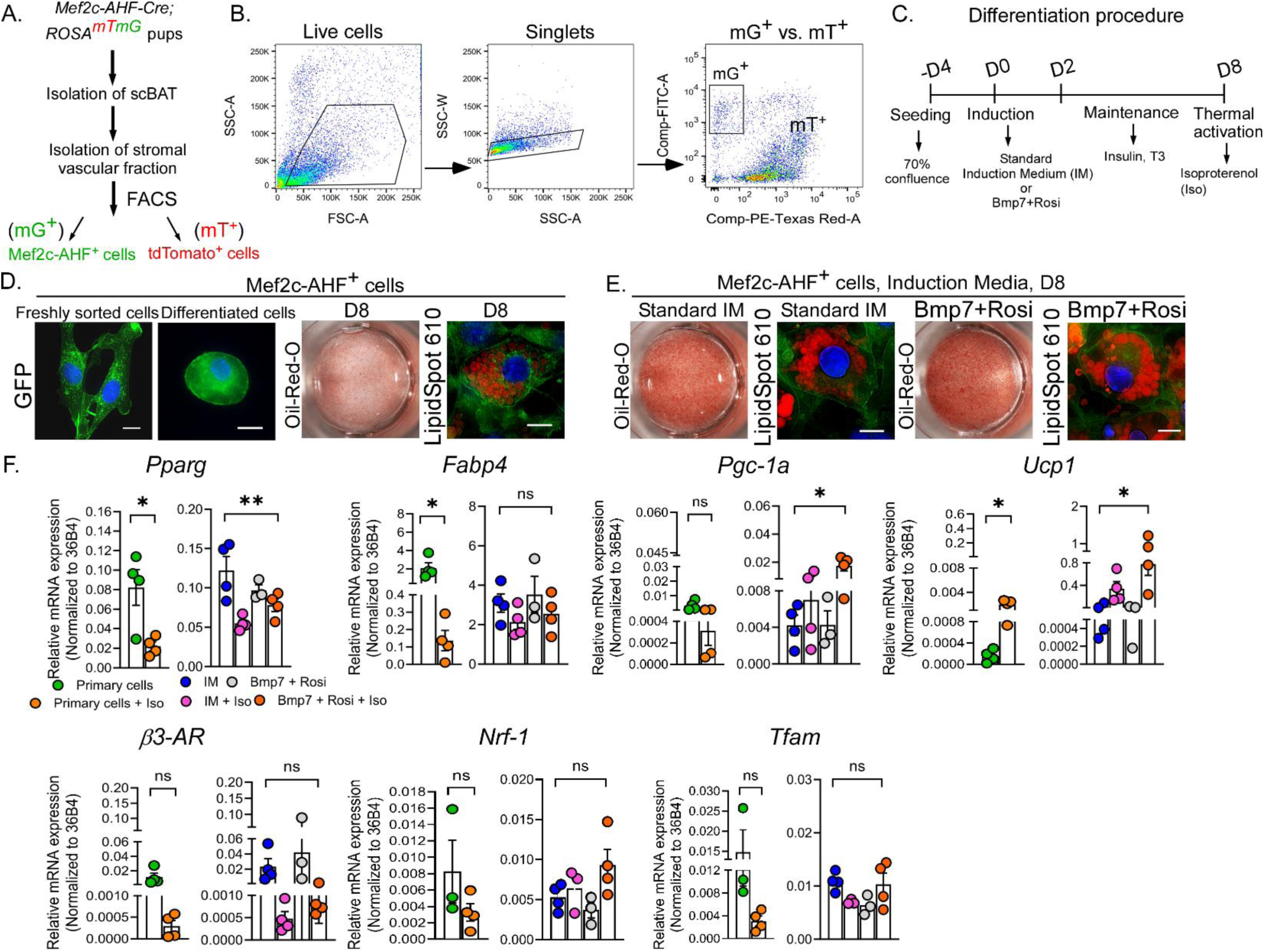
Mef2c-AHF-marked (Mef2c-AHF^+^) cells can differentiate into brown adipocytes. (A) An experimental design for isolating Mef2c-AHF^+^ cells from scBAT of postnatal *Mef2c-AHF- Cre;ROSA^mTmG^* mice. mG^+^: Mef2c-AHF^+^ cells. mT^+^: tdTomato^+^ cells. (B) Representative FACS plots of SVF isolated from the scBAT of postnatal 2.5- to 5-week-old *Mef2c-AHF-Cre;ROSA^mTmG^* mice. n = 28. (C) A schematic representation of the brown adipocyte differentiation procedure. Rosi: rosiglitazone. Iso: isoproterenol. (D) Representative images of freshly sorted and spontaneously differentiated Mef2c-AHF^+^ cells. The presence of lipid droplets in these cells is indicated by Oil-Red-O or LipidSpot 610. D8: day 8. Scale bar = 10 μm. n = 3-4. (E) Representative images of Mef2c-AHF^+^ cells treated with IM or Bmp7+Rosi. The presence of lipid droplets in these cells is indicated by Oil-Red-O or LipidSpot 610. Scale bar = 10 μm. n = 3-5. (F) Relative mRNA levels of genes involved in brown adipocyte differentiation and mitochondrial biogenesis in Mef2c-AHF^+^ cells that differentiated spontaneously or were induced by IM or Bmp7+Rosi. Iso was added to stimulate thermogenesis at the end of the culture period. Data are presented as mean ± SEM. **, p < 0.01; *, p < 0.05; ns = nonsignificant. n = 3-4.

Due to the limited number of sorted primary Mef2c-AHF^+^ cells, we generated an mG^+^ cell line (Mef2c-AHF-P5 cell) by immortalizing the SVF, followed by FACS to isolate mG^+^ cells from scBAT of the *Mef2c-AHF-Cre;ROSA^mTmG^* pups for more detailed molecular analyses (Figures S2A&2B). Although the immortalized mG^+^ cells lost their ability to spontaneously differentiate, they robustly differentiated into mature brown adipocytes upon IM induction, as indicated by high levels of Oil-Red-O-stained neutral lipids in these differentiated cells. RT-qPCR and western blotting confirmed the expression of the brown adipogenesis markers in these differentiated cells, specifically, the expression of UCP1 protein was highly induced in the differentiated cells (Figures S2C-E). Using this cell line, we also confirmed that Mef2c-AHF^+^ cells are Bmp pathway responsive. These cells express Bmp pathway receptors BMPR1A and BMPR2, and the transcription factor SMAD1, indicating they can be directly activated by the pathway ligand Bmp7 (Figure S2F). Indeed, Bmp7+Rosi treatment robustly induced the differentiation of Mef2c-AHF- P5 cells into mature brown adipocytes, as shown by the high level of Oil-Red-O-stained neutral lipids (Figure S2G) and the robust expression of key brown adipogenesis regulators in differentiated Mef2c-AHF-P5 cells (Figures S2H&I). Of note, the combined the use of IM and Bmp7+Rosi did not further enhance Mef2c-AHF-P5 differentiation (Figures S2H&I). Together, using both primary and immortalized Mef2c-AHF^+^ cells, we confirmed that Mef2c-AHF^+^ cells are progenitor cells that can give rise to supraclavicular brown adipocytes.

### IM and the combination of Bmp7 and Rosi induce distinctive gene expression profiles in early differentiating Mef2c-AHF^+^ cells

Both IM and Bmp7+Rosi can induce Mef2c-AHF^+^ cell differentiation, prompting us to investigate whether these two induction methods induce brown adipocyte differentiation similarly. We conducted RNA-sequencing (RNA-seq) to compare the expression profiles of untreated Mef2c-AHF-P5 cells to that of cells treated with IM, or Bmp7+Rosi for two days. Hierarchical clustering revealed that cells induced to differentiate by Bmp7+Rosi clustered together with untreated cells but were distinct from those induced by IM (Figure S3A). Notably, only 20% of the genes upregulated or downregulated by induction overlapped between IM- and Bmp7+Rosi-treated Mef2c-AHF-P5 cells (Figure S3B). Surprisingly, a disproportionately high number (70%) of genes were downregulated by IM but not by Bmp7+Rosi treatment (less than 10%) in these cells (Figure S3B). Gene ontology (GO) analysis further identified gene sets that are differentially regulated by both IM and Bmp7+Rosi or by IM or Bmp7+Rosi alone (Figures S3C&D). Lastly, heatmap clustering of genes involved in BAT-mediated thermogenesis further revealed that genes, such as *Ebf2*, *Zfp423*, and *Prdm16*, which are known to regulate early brown adipocyte differentiation, were more highly induced by Bmp7+Rosi than by IM (Figure S3E). To validate these RNA-seq results, we performed RT-qPCR to selectively examine the expression of genes that were preferentially induced by Bmp7+Rosi, such as *G0s2*, *Cd36*, and *Zfp423*, or by IM, such as *Lnc2* and *Dio2* (Figure S3F). Together, these analyses indicated that, while both IM and Bmp7+Rosi can robustly induce brown adipogenesis, the induction mechanism of Bmp7+Rosi is likely distinct from that of IM.

### Identifying cell surface markers expressed in Mef2c-AHF^+^ cells

Although lineage tracing can permanently mark Mef2c-AHF^+^ cells from scBAT, allowing isolation of these cells from genetically modified mice, identification of surface markers would allow for the isolation of brown adipocyte progenitors from non-genetically modified mice or other species, including human. Therefore, we performed flow cytometry to assess the potential for previously characterized cell surface markers expressed in the SVF of BAT or WAT to serve as markers for isolating Mef2c-AHF^+^ cells. Specifically, we focused on CD31, CD45, CD29, SCA-1, CD34, and CD24^36,37^. Additionally, markers known to be expressed in mesenchymal stem/progenitor cells, such as CD117, CD44, CD105, CD90, CD146, were included in the analysis^38^. Initially, the flow analysis was performed using SVF isolated from the scBAT of 5- to 7-day old *Mef2c-AHF-Cre;ROSA^mTmG^* pups. Due to the limited number of SVF cells obtained from newborn pups, these freshly isolated cells were cultured for 3 to 5 days before analysis. Among the analyzed markers, we found that the endothelial marker CD31 and immune cell marker CD45 were expressed at a very low level in the Mef2c-AHF^+^ cell (mG^+^) fraction (Figures 3A&B). Conversely, the stem cell markers CD29 and SCA-1 were highly expressed in the Mef2c-AHF^+^ cell fraction, while CD34 and CD24, markers for white preadipocytes, were expressed at low levels (Figures 3A&B). RT-qPCR confirmed the expression levels of these markers in sorted Mef2c-AHF^+^ cells (Figure 3C). We also found that mesenchymal stem cell markers (CD44, CD90, and CD146) were highly expressed, while CD105 was only moderately expressed in and CD117 was nearly absent from Mef2c-AHF^+^ cells (Figure S4A). As CD34 expression tends to be reduced in cultured cells^39,40^, we performed additional flow cytometry on the freshly isolated SVF from scBAT of 3- to 5-week-old Mef2c-*AHF-Cre;ROSA^mTmG^* mice to confirm this reduction in Mef2c-AHF^+^ cells. Again, a similar expression pattern was observed in these freshly sorted Mef2c-AHF^+^ cells (Figure S4B), and RT-qPCR verified the expression levels of these markers (Figure 4SC). Together, our flow cytometry analyses revealed dynamic expression of cell surface markers commonly found in the SVF of mouse adipose tissues in Mef2c-AHF^+^ cells.

**Figure 3.**
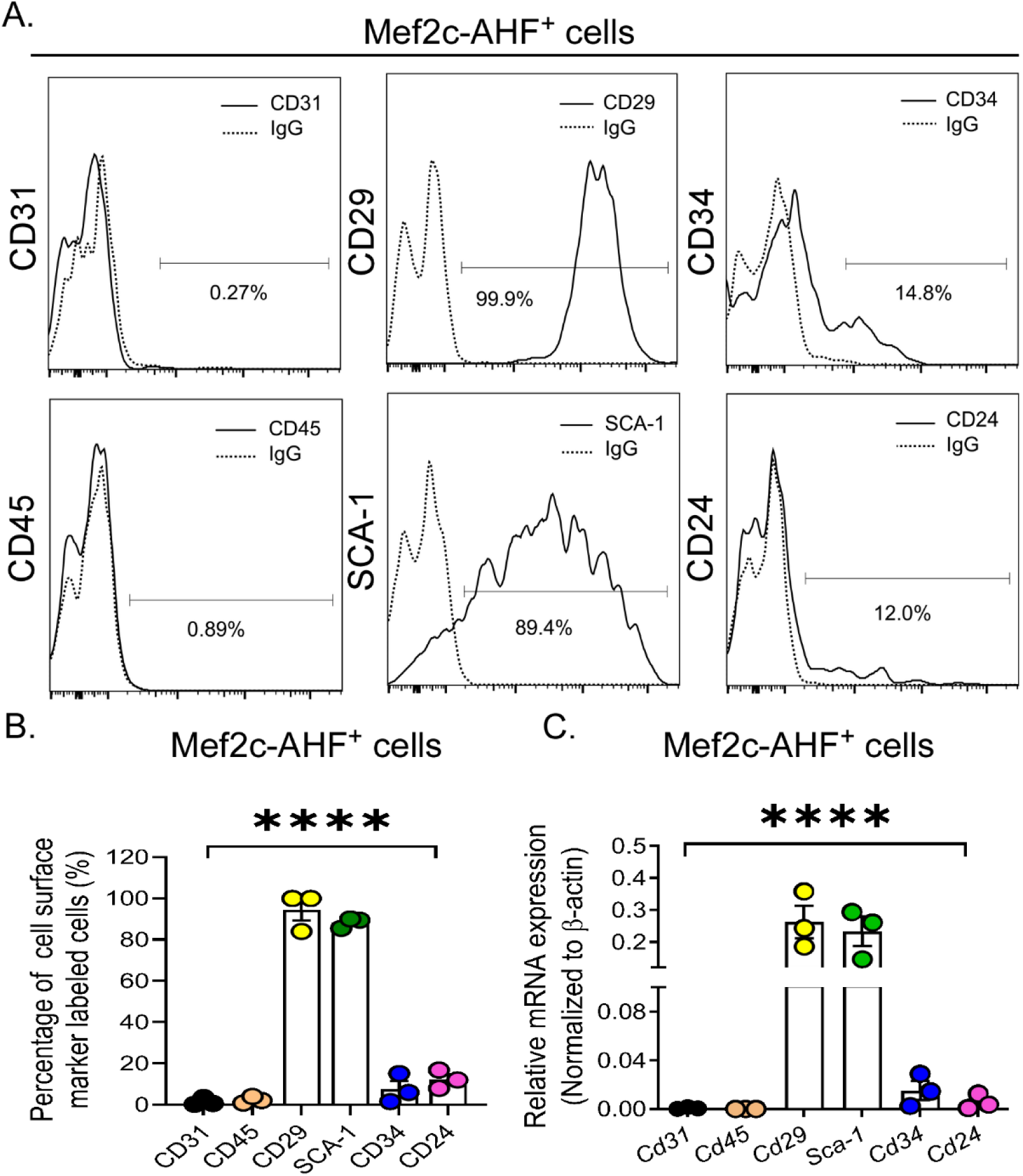
Flow cytometry analysis of cell surface marker expression in Mef2c-AHF^+^ cells. (A) Representative histogram plots showing the expression of CD31, CD45, CD29, SCA-1, CD34, and CD24, with the relevant isotype control (dashed line, IgG), in Mef2c-AHF^+^ cells. n = 3. (B) Quantification of the percentage of CD31, CD45, CD29, SCA-1, CD34, and CD24 positive cells in the Mef2c-AHF^+^ cell population compared to the relevant isotype control. Data are presented as mean ± SEM. ****, p < 0.0001. n = 3. (C) Relative mRNA levels of *Cd31*, *Cd45*, *Cd29*, *Sca-1*, *Cd34*, and *Cd24* in Mef2c-AHF^+^ cells isolated from the scBAT of postnatal 5- to 7-day-old *Mef2c-AHF-Cre;ROSA^mTmG^* mice. Data are presented as mean ± SEM. ****, p < 0.0001. n = 3.

**Figure 4.**
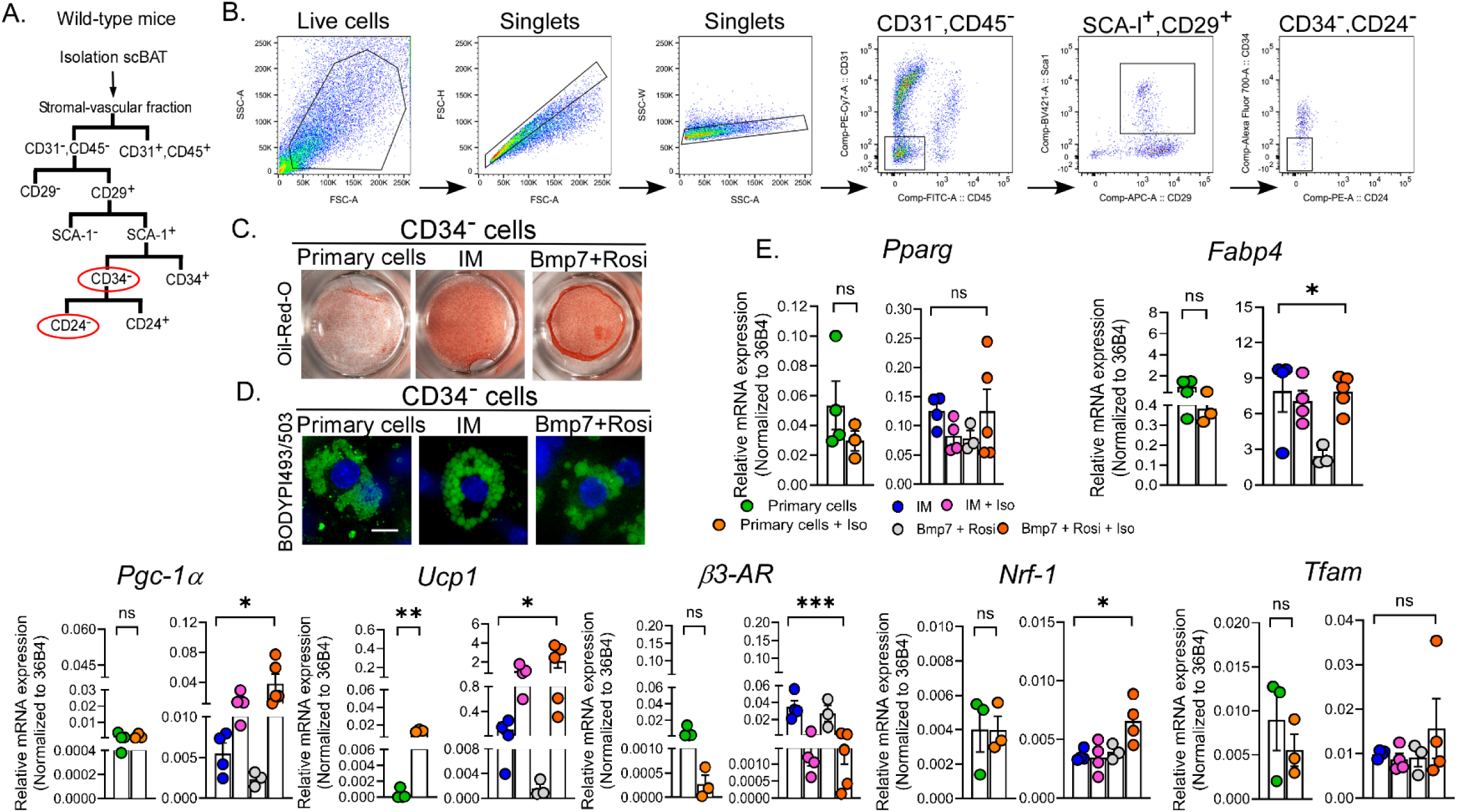
CD34^-^ cells isolated from the scBAT of wild-type mice can differentiate into brown adipocytes. (A) A schematic diagram of the FACS strategy for isolating CD34^-^ cells from scBAT of the postnatal wild-type mice. (B) Representative density plots showing FACS staining profiles and gating (black boxes) of SVF isolated from scBAT. n = 40. (C) Representative images of Oil-Red-O-stained CD34^-^ cells that either differentiated spontaneously or were induced by IM or by Bmp7+Rosi. n = 3. (D) Representative fluorescent images of BODIPY493/503-stained CD34^-^ cells that either differentiated spontaneously or were induced by IM or by Bmp7+Rosi. Scale bar = 10 μm. n = 3-4. (E) Relative mRNA levels of genes involved in brown adipocyte differentiation and mitochondrial biogenesis in CD34^-^ cells that differentiated spontaneously or were induced by IM or by Bmp7+Rosi. Iso was added to stimulate thermogenesis at the end of the culture period. Data are presented as mean ± SEM. ***, p < 0.001; **, p < 0.01; *, p < 0.05; ns = nonsignificant. n = 3-5.

### Isolation of [CD31^-^CD45^-^SCA-1^+^CD29^+^CD34^-^CD24^-^] (CD34^-^) cells from the scBAT of wild-type mice

To test the potential of using cell surface markers to isolate Mef2c-AHF-marked cells from scBAT, we established a panel of differentially expressed cell surface markers to use for FACS on the SVF isolated from scBAT of 2.5- to 5-week-old wild-type mice (Figure 4A). Both endothelial and immune cells were excluded from the SVF using CD31 and CD45. Cells highly expressing CD29 and SCA-1 were then purified from the CD31 and CD45 double-negative fraction. To further enhance purity, cells highly expressing CD34 and CD24 were excluded from the CD29 and SCA-1 double-positive fraction. We named this population of cells [CD31^-^CD45^-^SCA- 1^+^CD29^+^CD34^-^CD24^-^], abbreviated herein as CD34^-^ cells (Figure 4B). To verify that the CD34^-^ cells can give rise to brown adipocytes, like Mef2c-AHF^+^ cells do, we applied the same differentiation procedure listed in Figure 2C. The freshly sorted CD34^-^ cells were expanded once or twice before adipogenic induction. Like Mef2c-AHF^+^ cells, some CD34^-^ cells spontaneously differentiated into brown adipocytes during expansion (Figure 4C). CD34^-^ cells treated with IM or Bmp7+Rosi fully differentiated eight days after induction, as indicated by the abundance of Oil-Red-O-stained lipids (Figure 4C). BODIPY493/503 staining also confirmed the appearance of small lipid droplets in the differentiated CD34^-^ cells (Figure 4D). To verify that the differentiated CD34^-^ cells are brown adipocytes, we performed RT-qPCR to measure the expression of genes involved in brown adipogenesis, including *PPARg*, *Fabp4*, *Pgc-1α*, *Ucp1*, *beta3-AR*, *Nrf-1*, and *Tfam*. As shown in Figure 4E, all of these genes were expressed in CD34^-^ cells. Specifically, *Pgc- 1a* and *Ucp1* expression were significantly increased after Iso stimulation, indicating that the differentiated CD34^-^ cells are thermogenic brown adipocytes. These findings provide the first evidence that brown adipocyte stem/progenitor cells with properties similar to Mef2c-AHF^+^ cells can be isolated from wild-type scBAT. Thus, we present a new method to isolate these cells from non-genetically modified mice.

### Prrx1-marked progenitor cells give rise to supraclavicular brown adipocytes

After identifying the stem/progenitor cells that give rise to brown adipocytes in scBAT, we next investigated whether similar progenitor lineage relationships for scBAT exist in mouse and human. Earlier molecular profiling studies suggested that human scBAT has a molecular signature that is similar to that of mouse beige adipocytes in iWAT^3,24,25^. Since mouse scBAT does not originate from somite-derived cells like iBAT does, we hypothesized that mouse scBAT might share a lineage relationship with beige adipocytes in iWAT. To test this hypothesis, we performed lineage tracing using a Cre driver known to mark progenitor cells giving rise to iWAT, *Prrx1-Cre*.

We chose to trace Prrx1-marked cells because, like Mef2c-AHF-marked cells, they appear early during embryonic development^41,42^. To trace Prrx1-marked cells, we bred *Prrx1-Cre* mice with *R26R* mice to produce *Prrx1-Cre;R26R* progeny and performed X-gal staining. Consistent with early tracing studies, LacZ-positive cells were broadly expressed at E10.5 in *Prrx1-Cre;R26R* embryos (Figures S5A&B)^41^. To further trace Prrx1-marked cells, embryonic hearts were dissected at E12.5. At this stage, X-gal staining in AHF-derived regions of the heart largely overlapped with regions marked by Mef2c-AHF-Cre driver (Figure 5A). This overlap in the Cre-expressing domain suggests that Prrx1-marked cells, like Mef2c-AHF-marked cells, might give rise to scBAT. Indeed, X-gal-stained cells were present in scBAT but not iBAT from E18.5 *Prrx1-Cre;R26R* fetuses (Figure S5C). This observation was next confirmed in sectioned scBAT and iBAT from E18.5 *Prrx1-Cre;ROSA^mTmG^*fetuses and in postnatal 3-week-old mice (Figure 5B). Together, these tracing studies provide the first genetic evidence that scBAT and iWAT originate from the same progenitor cell population. To test if the Prrx1-marked (Prrx1^+^) cells can differentiate into brown adipocytes, we sorted these cells from the SVF isolated from scBAT of *Prrx1-Cre;ROSA^mTmG^* mice (Figures 5C&D). Prrx1^+^ cells isolated from the SVF of 2.5- to 5-week-old *Prrx1-Cre;ROSA^mTmG^*mice can differentiate spontaneously or be induced to differentiate by IM or Bmp7+Rosi, as shown by numerous Oil-Red-O- and LipidSpot 610-stained neutral lipids (Figure 5E). RT-qPCR also revealed that marker genes for brown adipogenesis were expressed in the sorted Prrx1^+^ cells (Figure 5F). As in Mef2c-AHF^+^ and CD34^-^ cells, Ucp1 expression was highly induced by Iso in the differentiated Prrx1^+^ cells (Figure 5F). Using genetic tracing and FACS, we provide new evidence that the Prrx1-Cre driver marks stem/progenitor cells that give rise to iWAT and scBAT.

**Figure 5.**
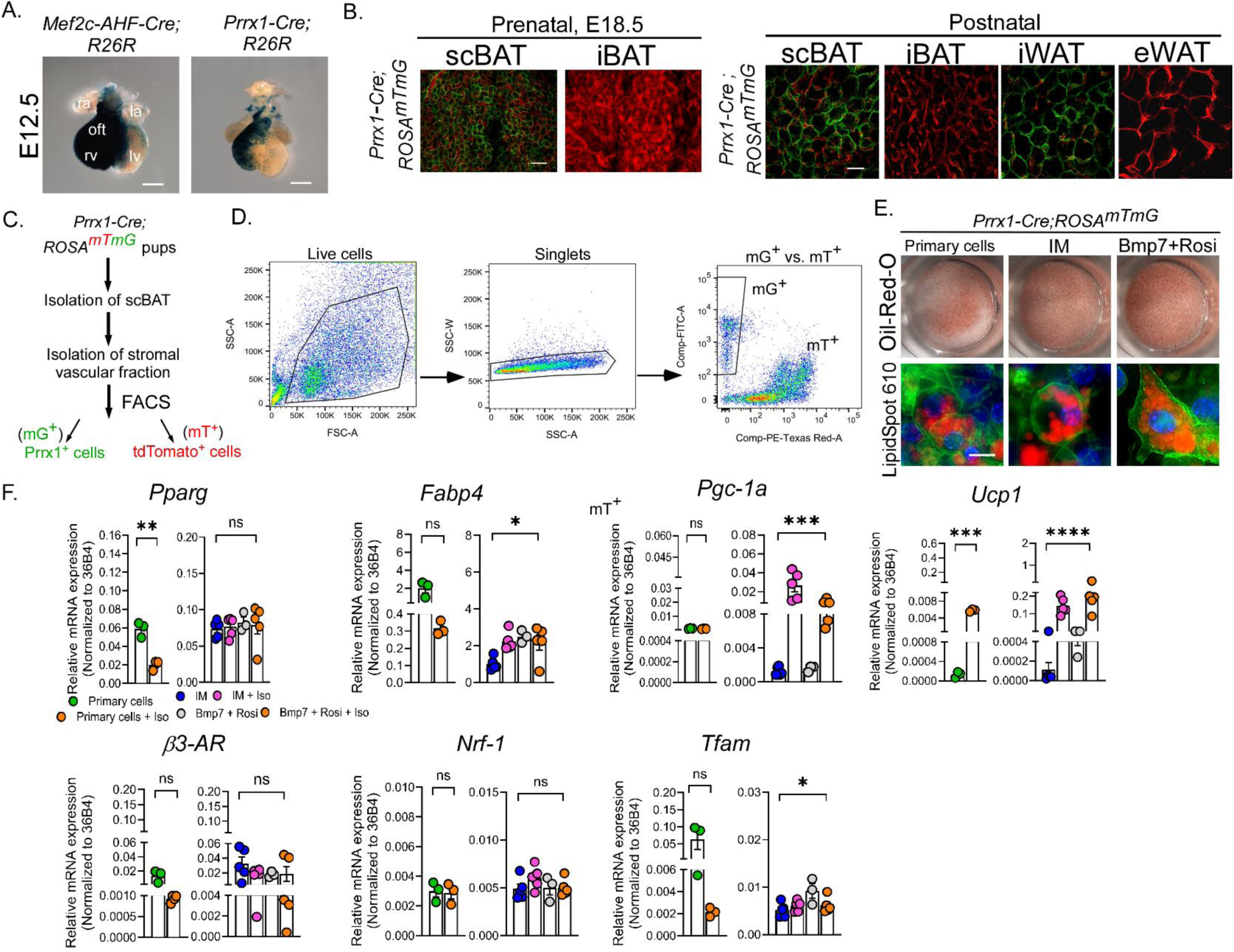
Prrx1 Cre Driver marks cells in both scBAT and iWAT. (A) Representative images of X-gal-stained hearts from E12.5 *Mef2c-AHF-Cre;R26R* and *Prrx1-Cre;R26R* embryos. Scale bar = 2000 μm. n = 4. ra: right atrium, rv: right ventricle, oft: outflow tract, la: left atrium, lv: left ventricle. (B) Representative fluorescence images of scBAT and iBAT from E18.5 and postnatal *Prrx1-Cre;ROSA^mTmG^*mice. Scale bar = 25 μm. n = 3-4. (C) Experimental design for isolating Prrx1^+^ cells from the scBAT of postnatal *Prrx1-Cre;ROSA^mTmG^* mice. mG^+^: Prrx1^+^ cells. mT^+^: tdTomato^+^ cells. (D) Representative FACS plots of SVF isolated from scBAT of the postnatal *Prrx1-Cre;ROSA^mTmG^* mice. n = 20. (E) Representative images of Oil-Red-O and LipidSpot 610-stained Prrx1^+^ cells that differentiated spontaneously or were induced by IM or by Bmp7+Rosi. Scale bar: 10 μm. n = 4-5. (F) Relative mRNA levels of genes involved in brown adipocyte differentiation and mitochondrial biogenesis in Prrx1^+^ cells that differentiated spontaneously or were induced by IM or by Bmp7+Rosi. Iso was added to stimulate thermogenesis at the end of the culture period. Data are presented as mean ± SEM. ****, p < 0.0001; ***, p < 0.001; **, p < 0.01; *, p < 0.05; ns = nonsignificant. n = 3-5.

### Isolation of CD34^-^ cells from iWAT of the wild-type mice

Although tracing of Prrx1-marked cells established a lineage relationship between scBAT and iWAT, the Prrx1 Cre driver marks both beige and white adipocytes in iWAT, preventing direct examination of cellular similarity between brown adipocytes from scBAT and beige adipocytes from iWAT. Therefore, we applied the cell surface marker panel for sorting CD34^-^ cells from scBAT and isolated a group of CD34^-^ cells from the iWAT of 3-5-week-old wild-type mice (Figures 6A&B). To test the differentiation potential of these CD34^-^ cells from iWAT, we applied a widely accepted beige adipocyte differentiation procedure, incorporating Rosi and triiodothyronine (T3) into the IM culture medium (Figure 6C). To increase cell numbers, we expanded the sorted CD34^-^ cells once or twice before induction. Surprisingly, like CD34^-^ cells from scBAT, those from iWAT slowly and spontaneously differentiated during expansion. Their differentiation efficiency was further enhanced by IM or by a combination of IM, Rosi, and T3. Again, Oil-Red-O staining was applied to reveal the appearance of lipid-filled CD34^-^ cells (Figure 6D). BODIPY493/503 staining further confirmed numerous small lipid droplets in the differentiated CD34^-^ cells (Figure 6E). RT-qPCR verified the expression of marker genes for beige adipogenesis (Figure 6F). Overall, the CD34^-^ cells from iWAT exhibit a similar expression profile to those from scBAT. Specifically, *Ucp1* expression and UCP1-mediated uncoupling were much higher in CD34^-^ cells receiving the beige adipocyte induction cocktail compared to those treated with IM (Figure 6G). Together, this molecular evidence indicates that CD34^-^ cells from iWAT show significant similarities to those from scBAT. These cellular similarities support our genetic tracing studies, which demonstrate that stem/progenitor cells, descending from the same progenitor cells marked by the Prrx1-Cre driver, give rise to both brown adipocytes of scBAT and beige adipocytes of iWAT.

**Figure 6.**
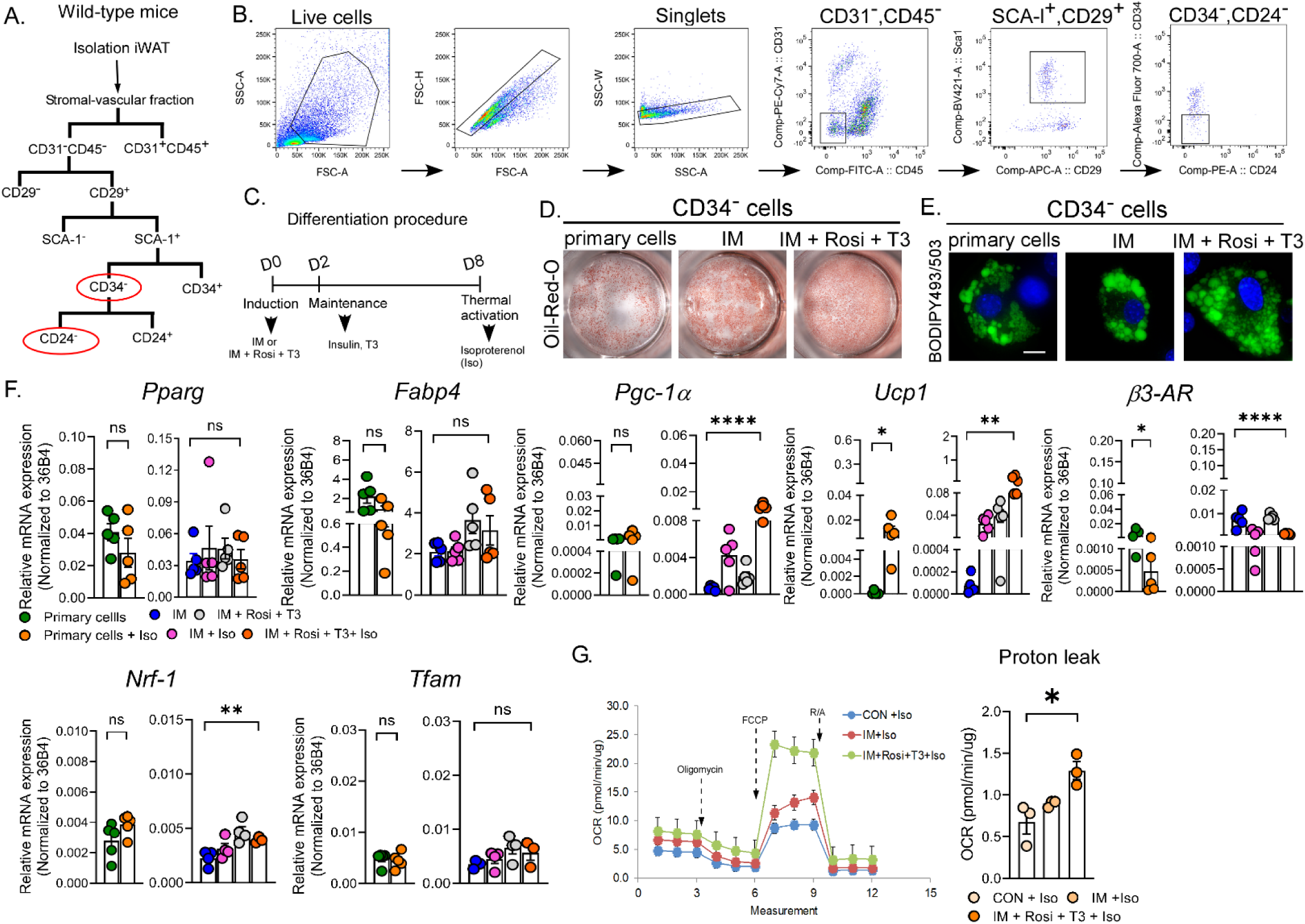
CD34^-^ cells isolated from the iWAT of wild-type mice can differentiate into beige adipocytes. (A) A schematic diagram of the FACS strategy for isolating CD34^-^ cells from the iWAT of postnatal wild-type mice. (B) Representative density plots showing FACS staining profiles and gating (black boxes) of SVF isolated from iWAT. n = 58. (C) A schematic representation of the beige adipocyte differentiation procedure. (D) Representative images of Oil-Red-O-stained CD34^-^ cells that differentiated spontaneously or were induced by IM or by a combination of IM, Rosi, and T3. n = 3-4. (E) Representative fluorescent images of BODIPY493/503-stained CD34^-^ cells that differentiated spontaneously or were induced by IM or by a combination of IM, Rosi, and T3. Scale bar = 10 μm. n = 4-6. (F) Relative mRNA levels of genes involved in beige adipogenesis and mitochondrial biogenesis in CD34^-^ cells that differentiated spontaneously or were induced by IM or by a combination of IM, Rosi, and T3. Iso was added to stimulate thermogenesis at the end of the culture period. Data are presented as mean ± SEM. ****, p < 0.0001; **, p < 0.01; *, p < 0.05; ns = nonsignificant. n = 3-5. (G) Representative Seahorse assay traces and a graph of calculated protein leak (uncoupling) in CD34^-^ cells that differentiated spontaneously or were induced by IM or by a combination of IM, Rosi, and T3. Iso was added to stimulate thermogenesis at the end of the culture period. Oxygen consumption rate (OCR) values were normalized to the total protein amount. Data are presented as mean ± SEM. *, p < 0.05. n = 3.

Given that previous studies have shown that white adipocytes originate from progenitor cells expressing CD34^36^, in this study, we also examined the cellular properties of [CD31^-^CD45^-^ SCA-1^+^CD29^+^CD34^+^CD24^-^] cells, referred to as CD34^+^ cells. We isolated the CD34^+^ fraction from the SCA-1 and CD29 double-positive fraction (Figures S6A&B). Again, we expanded the sorted CD34^+^ cells once or twice before adipogenic induction. Like CD34^-^ cells, CD34^+^ cells gradually differentiated during cell expansion (Figure S6C). The differentiation efficiency of these cells was further enhanced by IM or a combination of IM, Rosi, and T3, as shown by increased Oil-Red-O-stained CD34^+^ cells (Figure S6C). Unlike CD34^-^ cells, BODIPY493/503 staining revealed large lipid droplets in the differentiated CD34^+^ cells, characteristic of white adipocytes (Figure S6D). RT-qPCR revealed the markers for adipogenesis, *Pparg* and *Fabp4*, were expressed in CD34^+^ cells, but thermogenic markers *Pgc-1α* and *Ucp1* increased less than in CD34^-^ cells (Figure S6E). UCP1-mediated uncoupling was also significantly lower in CD34^+^ cells than in CD34^-^ cells (Figure S6F). Overall, our data suggest that CD34^+^ cells give rise to adipocytes resembling white adipocytes, with much lower thermogenic potential.

### Progenitors of scBAT could originate from the early developing heart

In this study, we found that AHF-derived cells marked by *Mef2c-AHF-Cre* or *Prrx1-Cre* mice, which are known to pattern parts of the developing heart, contribute to the formation of scBAT. Since the heart develops earlier than scBAT, we reasoned that scBAT progenitors could be within the developing heart and marked by Mef2c-AHF- and Prrx1-Cre drivers. To test this possibility, we developed an intersectional tracing system for precise tracing of the cell population that is marked by Mef2c-AHF and Prrx1 in the developing heart. This tracing system uses dual-recombinase (a Cre and a Dre deriver) and an intersectional ROSA^mTmG^ (*iROSA^mTmG^*) reporter mouse. We generated the *iROSA^mTmG^* by knocking a targeting cassette containing a membrane-targeted tdTomato (mT) and a membrane-targeted GFP (mG) into the *ROSA26* locus (Figures S7A&B). Under this design, mT is activated upon Cre-mediated excision, while mG is activated upon Cre and Dre-mediated excision. The cell populations marked by Cre alone or by both Cre and Dre can be separately traced (Figure S7C). To trace progenitors of scBAT, we bred the *Prrx1-Cre;Mef2c-AHF-DreERT2* double heterozygous male mice with the *iROSA^mTmG^* female mice (Figure 7A). Dre was activated by injecting Tam at E6.5, prior to the appearance of the Mef2c-AHF-marked cells and analyzed at E9.5 (Figure 7B). As shown in Figure 7C, Prrx1-marked (mT) cells were expressed in the midbrain, forelimb, outflow tract, and right ventricle of the developing heart at E9.5. Unexpectedly, we also observed a group of Mef2c-AHF and Prrx1-dual marked (mG^+^) cells in the outflow tract and right ventricle (Figure 7C). This observation was also noted in embryos that received Tam injection at E8.5 and were analyzed at E10.5 and E12.5. The mG^+^ cells appeared to be concentrated in the outflow tract (Figures 7B-D). To further determine whether these mG^+^ cells are involved in scBAT formation, Tam was injected at E12.5, before the scBAT primordium developed in the embryo’s neck, and analyzed at E16.5. As shown in Figures 7B&E, mG^+^ cells were highly abundant in the primordium of scBAT at E16.5. In summary, using a dual recombinase-mediated intersectional tracing system, we uncovered that the progenitors of scBAT may emerge early during cardiac development.

**Figure 7.**
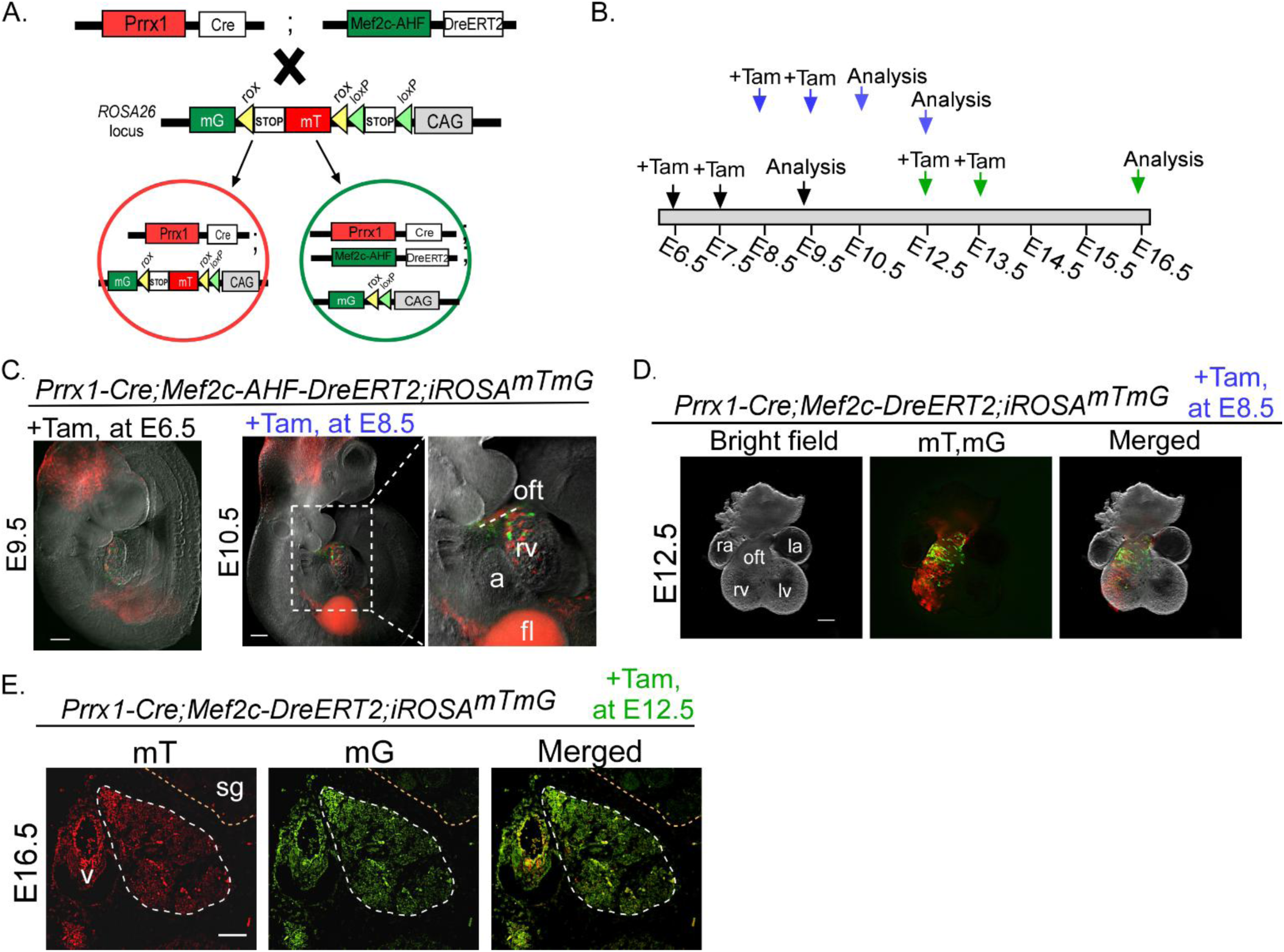
The progenitors of scBAT originate from early developing heart. (A) A schematic diagram of the intersectional lineage tracing strategy using *Prrx1-Cre* and *Mef2c-AHF-DreERT2* mice crossed with intersectional ROSA^mTmG^ (*iROSA^mTmG^*) mice to identify progenitors of scBAT. (B) A schematic graph outlining the timing and frequencies of the tamoxifen (Tam) treatments to induce Dre expression. (C) Representative images of E9.5 and E10.5 *Prrx1-Cre;Mef2c-AHF- DreERT2;iROSA^mTmG^* embryos. Cells expressing mT are Prrx1^+^ cells, and cells expressing mG are Prrx1^+^ and Mef2c-AHF^+^ cells. Scale bar = 2500 μm. n = 3-4. a: atrium, fl: forelimb, rv: right ventricle, oft: outflow tract. (D) Representative images of heart from E12.5 *Prrx1-Cre;Mef2c- AHF-DreERT2;iROSA^mTmG^* embryos. Scale bar = 2500 μm. n = 5. (E) Representative images of scBAT from E16.5 *Prrx1-Cre;Mef2c-AHF-DreERT2;iROSA^mTmG^* embryos. Scale bar = 100 μm. n = 2. v: vein. sg: salivary gland.

## Discussion

While the importance of BAT in metabolism has been recognized in mice and humans, the origin of BAT, specifically, the scBAT remains largely unexplored. Using single and intersectional genetic tracing, we discovered that many brown adipocytes in scBAT descend from Mef2c and Prrx1-marked AHF progenitor cells.

The discovery that scBAT is derived from progenitor cells that also contribute to the heart, is unexpected because adipocytes are not typically found in a healthy developing heart. Since it was generated two decades ago, the *Mef2c-AHF-Cre* mouse has been widely used in genetic and functional studies of cardiac tissues-derived from AHF. Recent clonal expression analyses indicated that Mef2c-AHF-marked cells can give rise to cranial and neck muscles and veins, suggesting a shared origin of these cells with the developing heart^33^. Here, our tracing data adds brown adipocytes to the increasing list of cell types descending from Mef2c-AHF-marked cells outside the heart. Identifying other cell types, besides adipocyte stem/progenitors, that also descend from Mef2c-AHF-marked cells in scBAT would be of great interest to researchers for understanding complex organ development and the lineage relationships among cells in those organs.

Another unexpected observation from this study is that sorted Mef2c-AHF^+^ cells spontaneously differentiated into brown adipocytes in the growth medium provided. This persistent phenomenon suggests that Mef2c-AHF^+^ cells likely possess stem cell properties. Indeed, our cell surface marker analysis revealed that Mef2c-AHF^+^ cells highly express cell surface markers, such as SCA-1, CD29, CD44, CD90, and CD146, for mouse and human mesenchymal stem cells. These analyses, thus, support our *in vitro* observation that Mef2c-AHF^+^ cells are a group of previously unknown adipocyte stem cells. Remarkably, the ability of adipocyte stem/progenitor cells to differentiate spontaneously *in vitro* is not limited to brown adipocyte stem/progenitor cells. As reported earlier, a group of LY6C^−^CD9^−^PDGFRβ^+^ cells isolated from eWAT can also differentiate under a culture medium similar to the one used in this report ^43^. Lastly, while lineage tracing allows for permanent marking of adipocyte stem/progenitor cells *in vivo*, it requires time-consuming generation of genetically modified mice. Importantly, we developed a panel of cell surface markers expressed in Mef2c-AHF^+^ cells to isolate CD34^-^ cells with similar cellular and molecular properties. Using this panel of cell surface markers, it should be possible to isolate adipocyte stem/progenitor cells from wild-type mice or even human BAT for future translational studies.

Since the discovery of mouse scBAT, it has been our intention to probe the lineage origins of both mouse scBAT and human BAT. Mouse scBAT is situated in the neck where the most visible human BAT is also found. Although genetic tracing is not feasible for human studies, molecular profiling suggested that molecular signatures for human BAT are more similar to mouse beige adipocytes, despite morphological similarities between human and mouse BAT. In this report, we provide genetic evidence that mouse scBAT is more closely related to iWAT than iBAT, as indicated by *Prrx1-Cre* mice marking both scBAT and iWAT, but not iBAT. Because iWAT is composed of both white and beige adipocytes, single lineage tracing cannot separately trace Prrx1^+^ white and beige adipocytes in iWAT. Alternatively, we used cell surface markers, initially applied to sort CD34^-^ cells from the SVF of scBAT, to similarly sort CD34^-^ cells from the SVF of iWAT. As shown in this report, the cellular and molecular properties of CD34^-^ cells isolated from iWAT resemble those of scBAT but are distinct from those of CD34^+^ cells from the SVF of iWAT. Similarly, a group of thermogenic CD34^-^ cells was recently isolated from human WAT^44^. Altogether, these cell-based studies provide *in vitro* evidence that brown adipocytes in scBAT and beige adipocytes share similar molecular properties, which could have translational implications for further understanding human brown adipose biology.

Although it was not previously clear at which developmental stage human BAT depots, including scBAT, begin to form, our studies indicate that mouse scBAT develops prenatally. To identify the progenitors of scBAT, we developed a new intersectional tracing system. Using this system, we traced a group of stem/progenitor cells marked by both Mef2c-AHF-and Prrx1-Cre drivers that appeared in the developing heart as early as E9.5. Thus, despite forming at later developmental stages, progenitors of scBAT seem to appear early during development. Similar results were also observed in WAT. Progenitor cells that give rise to white adipocytes were also found to emerge around E10.5 in mice^45,46^. While much of the current research in the field focuses on understanding the mechanisms underlying the differentiation of preadipocytes into mature adipocytes, the lineage trajectory and mechanisms that direct adipocyte stem/progenitors to differentiate toward preadipocytes should be further explored to gain a full understanding of brown and white adipogenesis.

### Limitations of study

Both Mef2c-AHF and Prrx1-marked progenitor cells emerge early during development and can give rise to multiple cell types in the heart and neck prior to the formation of scBAT. It is not yet clear whether these cells directly give rise to scBAT or this process involves additional intermediate cell types. A recent study reported that a subset of scBAT could originate from Tbx1- marked myogenic progenitor cells^47^. We could not address this complex developmental process in this paper with currently available lineage tracing mice. However, future studies using a dual-recombinase-mediated intersectional lineage tracing system will enable further dissection of the specific subsets of cells (myogenic or non-myogenic) descending from Mef2c-AHF or Prrx1-marked progenitors that contribute to scBAT. In this report, we employed genetic lineage tracing approaches to first trace the origin of scBAT. While this method allowed us to permanently mark stem/progenitor cells that give rise to scBAT, it cannot reveal the transcriptional network governing the development of scBAT. Future studies using bulk- and single cell RNA-seq will reveal specific transcriptional signatures of Mef2c-AHF and Prrx1-marked cells, allowing for mechanistic studies. Lastly, our studies revealed that Mef2c-AHF and Prrx1-marked cells contribute to scBAT during development. Whether these cells also contribute to the generation of adult scBAT will need to be defined in future studies.

## Acknowledgments

The authors wish to thank Mrs. Yufeng Shi for assisting in mouse husbandry and gene expression analysis, Mrs. Tatiana Goltsova and staff at the Texas Children’s Hospital Flow Cytometry Core Laboratory for performing flow cytometry and FACS, staff members at the BCM Genetically Engineered Mouse Core for rederiving *Mef2c-AHF-Cre* and *CAG-Dre* mice, and Ms. Gladys Zapata and Mr. Nitesh Mehta at the Laboratory for Translational Genomics for RNA sequencing. The authors also wish to thank Dr. Benoit Bruneau (Gladstone Institutes, San Francisco, CA) for providing *Mef2c-AHF-DreERT2* and *ROSA^nKmB^* mice, and Dr. Tong Qiang for providing HEK293T cells. This study was supported by NIH NIDDK R01DK116899; the American Heart Association grant 16GRNT30720003; a pilot award from the Baylor College of Medicine Cardiovascular Research Institute to M.H.C.; NIH grants HL177462 and NS126499 to B.L.B.; NIH NCI R00 CA237618 to X.G.; USDA/ARS CRIS grant 3092-51000-064-05S to M.H.C and X.G, and USDA/ARS grant CRIS 3092-51000-065-003S and the Research Vision at Texas Children’s Hospital to H.K.Y.

## Author contributions

Conceptualization: M.H.C. B.L.B.; Methodology: M.H.C., Y.R. Y.T.S.; Validation: M.H.C., Y.R., K.Z., Z.W.; Formal analysis: M.H.C., Y.R., K.Z., Y.T.S., Z.W.; Investigation: M.H.C., Y.R., K. Z., Y.T.S., Q.M., Z.W., H.K.Y., S.J., M.K., Z.G., C.W., T.S.; Data curation: Q.M., Z.W. M.H.C., H. K.Y.; Writing - original draft: M.H.C., Y.R., Y.T.S., S.J.; Writing - review & editing: M.H.C., Y.R., K.Z., Y.T.S., Q.M., Z.W., H. K.Y., S.J., M.K., X.G., C.W., T.S., B.L.B.

## Supplemental Figure Legends

**Figure S1.**
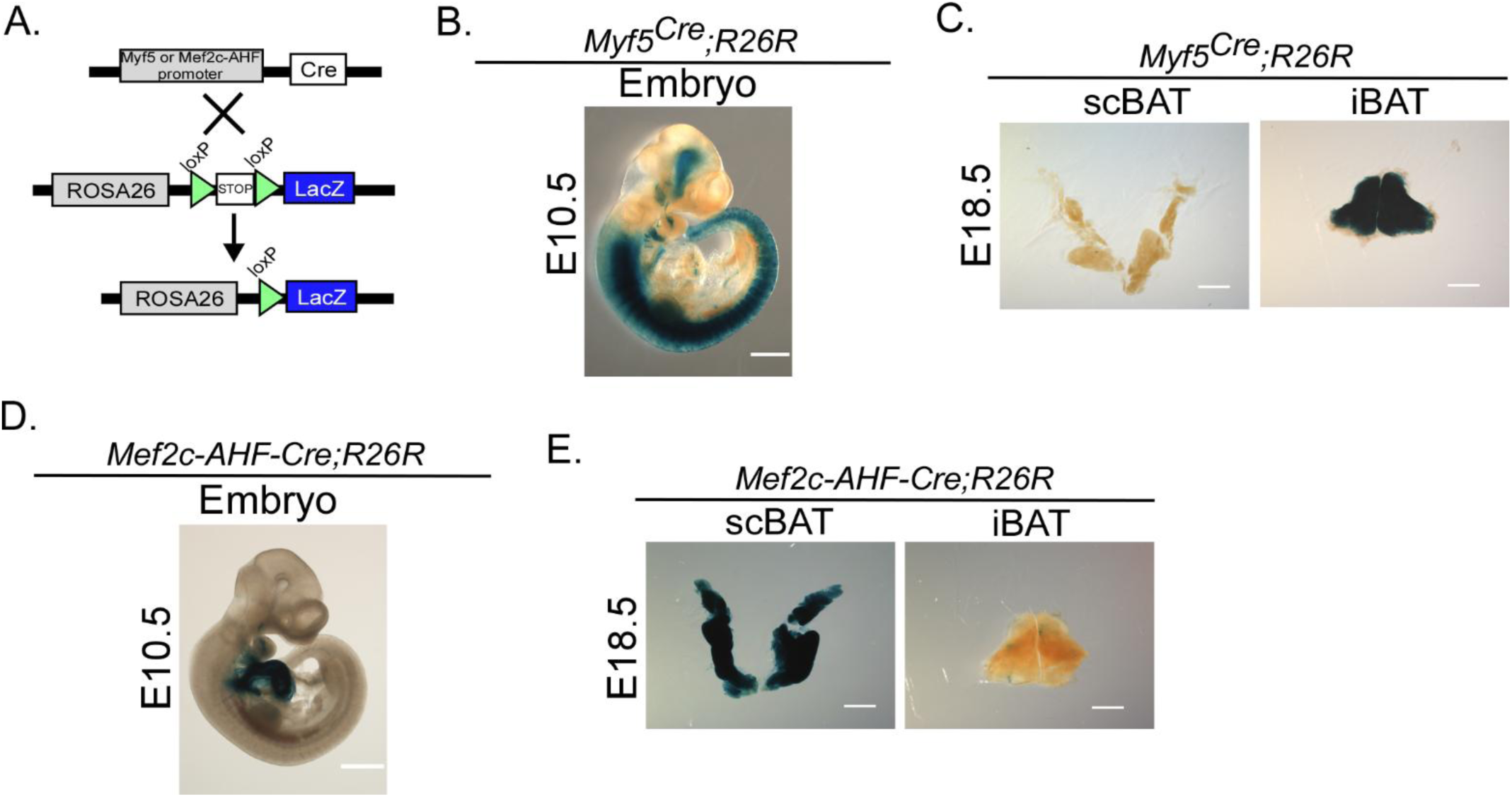
A Myf5^Cre^ driver marks iBAT, while a Mef2c-AHF-Cre driver marks scBAT in mouse embryos. (A) A schematic diagram of the lineage tracing strategy using *Myf5^Cre^* or *Mef2c- AHF-Cre* mice crossed with *R26R* mice to identify Myf5- or Mef2c-AHF-marked BAT. (B) Representative image of whole mount X-gal-stained embryos from E10.5 *Myf5^Cre^;R26R* embryos. Scale bar = 2000 μm. n = 5. (C) Representative images of X-gal-stained scBAT and iBAT from E18.5 *Myf5^Cre^;R26R* embryos. Scale bar = 2000 μm. n = 3. (D) Representative image of whole mount X-gal-stained embryos from E10.5 *Mef2c-AHF-Cre;R26R* embryos. Scale bar = 2000 μm. n = 4. (E) Representative images of X-gal-stained scBAT and iBAT from E18.5 *Mef2c-AHF- Cre;R26R* embryos. Scale bar = 2000 μm. n = 4.

**Figure S2.**
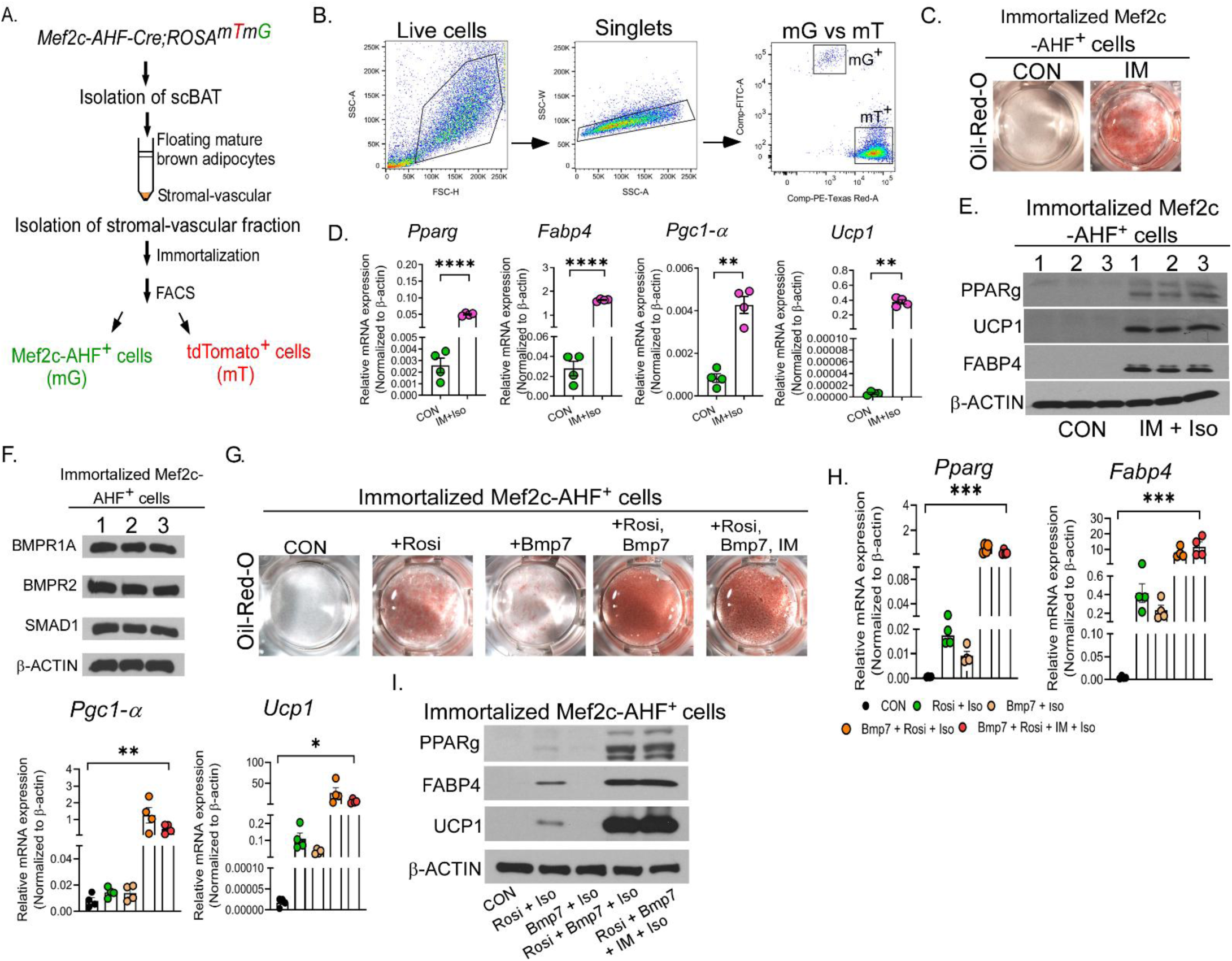
Immortalized Mef2c-AHF^+^ (Mef2c-AHF-P5) cells can be induced to differentiate into mature brown adipocytes by IM or by Bmp7+Rosi. (A) A schematic diagram of the FACS strategy for isolating immortalized Mef2c-AHF^+^ cells. (B) Representative FACS plots of immortalized SVF isolated from the scBAT of postnatal-day-5 (P5) *Mef2c-AHF-Cre;ROSA^mTmG^* mice. (C) Representative images of Oil-Red-O-stained Mef2c-AHF-P5 cells undergoing differentiation induced by IM. n = 3-4. CON: non-differentiated. (D) Relative mRNA levels of genes involved in brown adipogenesis in CON and IM-treated Mef2c-AHF-P5 cells. Iso was added to stimulate thermogenesis at the end of the culture period. Data are presented as mean ± SEM. ****, p < 0.0001; **, p < 0.01. Iso: Isoproterenol. n = 4. (E) Relative protein levels of PPARg, UCP1, and FABP4 in CON and IM-treated Mef2c-AHF-P5 cells. Iso was added at the end of the culture period. n = 3. β-ACTIN is used as a loading control. (F) Western blot analysis of the components of the Bmp signaling pathway, including receptors, BMPR1A and BMPR2, and the transcription factor SMAD1, in Mef2c-AHF-P5 cells. β-ACTIN is used as a loading control. n = 3. (G) Representative images of Oil-Red-O-stained Mef2c-AHF-P5 cells undergoing differentiation induced by Rosi, Bmp7, Bmp7+Rosi, or a combination of Bmp7+Rosi and IM. n = 3. (H) Relative mRNA levels of genes involved in brown adipogenesis in CON, Rosi, Bmp7, Bmp7+Rosi, or a combination of Bmp7+Rosi and IM-treated Mef2c-AHF-P5 cells. Iso was added at the end of the culture period. Data are presented as mean ± SEM. ***, p < 0.001; **, p < 0.01; *, p < 0.05. n = 4. (I) Relative protein levels of PPARg, UCP1, FABP4 in CON, Rosi, Bmp7, Bmp7+Rosi, or a combination of Bmp7+Rosi, and IM-treated Mef2c-AHF-P5 cells. Iso was added at the end of the culture period. n = 2. β-ACTIN is used as a loading control.

**Figure S3.**
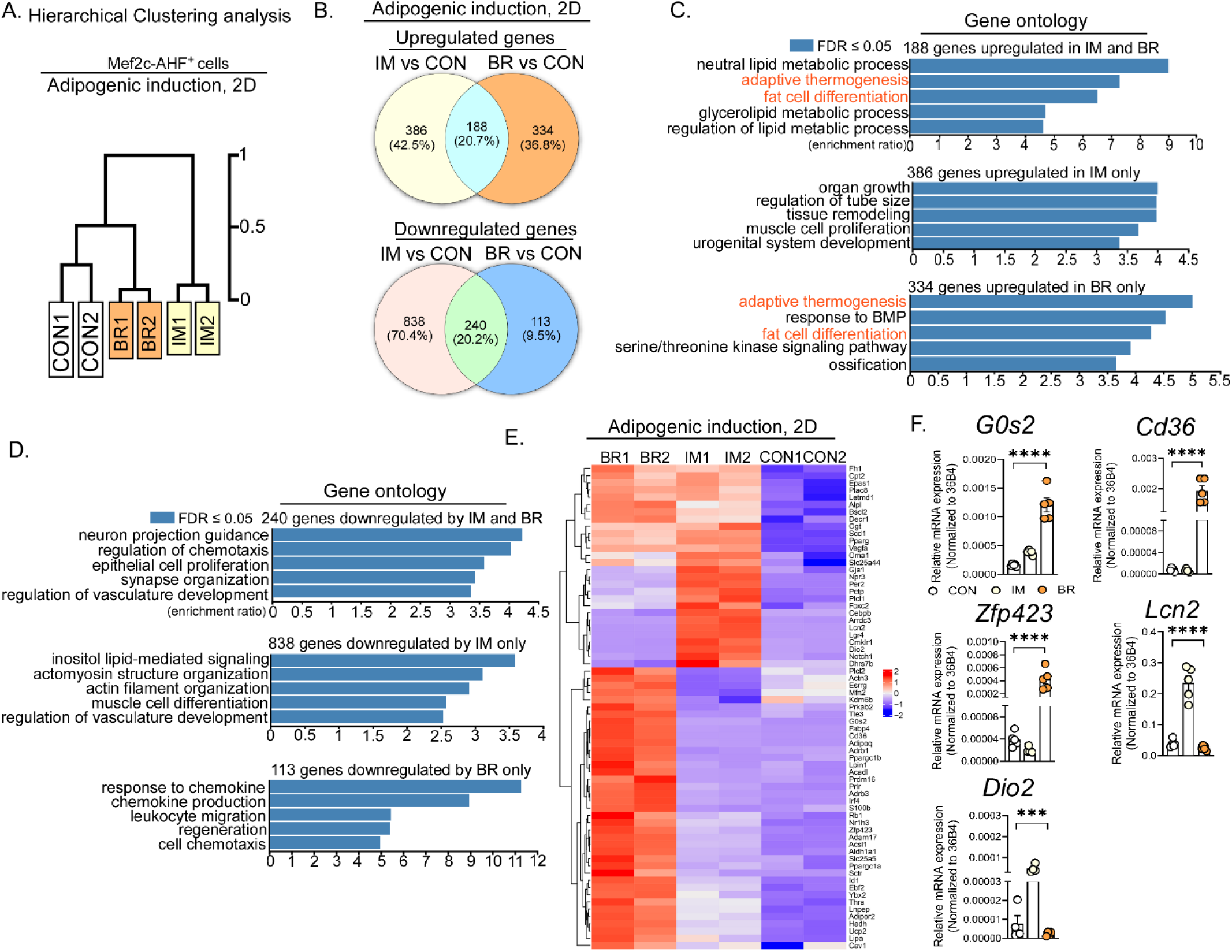
Comparative RNA-seq analysis reveals differentially expressed genes in Mef2c- AHF-P5 cells induced to differentiate by IM or by a combination of Rosi and Bmp7 for 2 days. (A) Hierarchical clustering of the genes in non-differentiated (CON), IM, or Bmp7+Rosi (BR)-induced Mef2c-AHF-P5 cells. (B) Venn diagrams showing the number of upregulated and downregulated genes in CON, IM, or Bmp7+Rosi-induced Mef2c-AHF-P5 cells. Genes with expression that differed by more than twofold were used in the analysis. (C & D) Top 5 Gene Ontology (GO) terms of the upregulated gene sets (C) and downregulated gene sets (D) shown in B. (E) Hierarchical clustering heat map of the differentially expressed genes involved in thermogenesis in CON, IM, or Bmp7+Rosi-induced Mef2c-AHF-P5 cells. (Red = upregulated compared to CON. Blue = downregulated compared to CON.). (F) RT-qPCR verification of the genes identified by RNA-seq, including *G0s2*, *Cd36*, *Zfp423*, *Lcn2*, and *Dio2*, as being upregulated in IM or Bmp7+Rosi-induced Mef2c-AHF-P5 cells. Data are presented as mean ± SEM. ****, p < 0.0001; ***, p < 0.001. n = 4-5.

**Figure S4.**
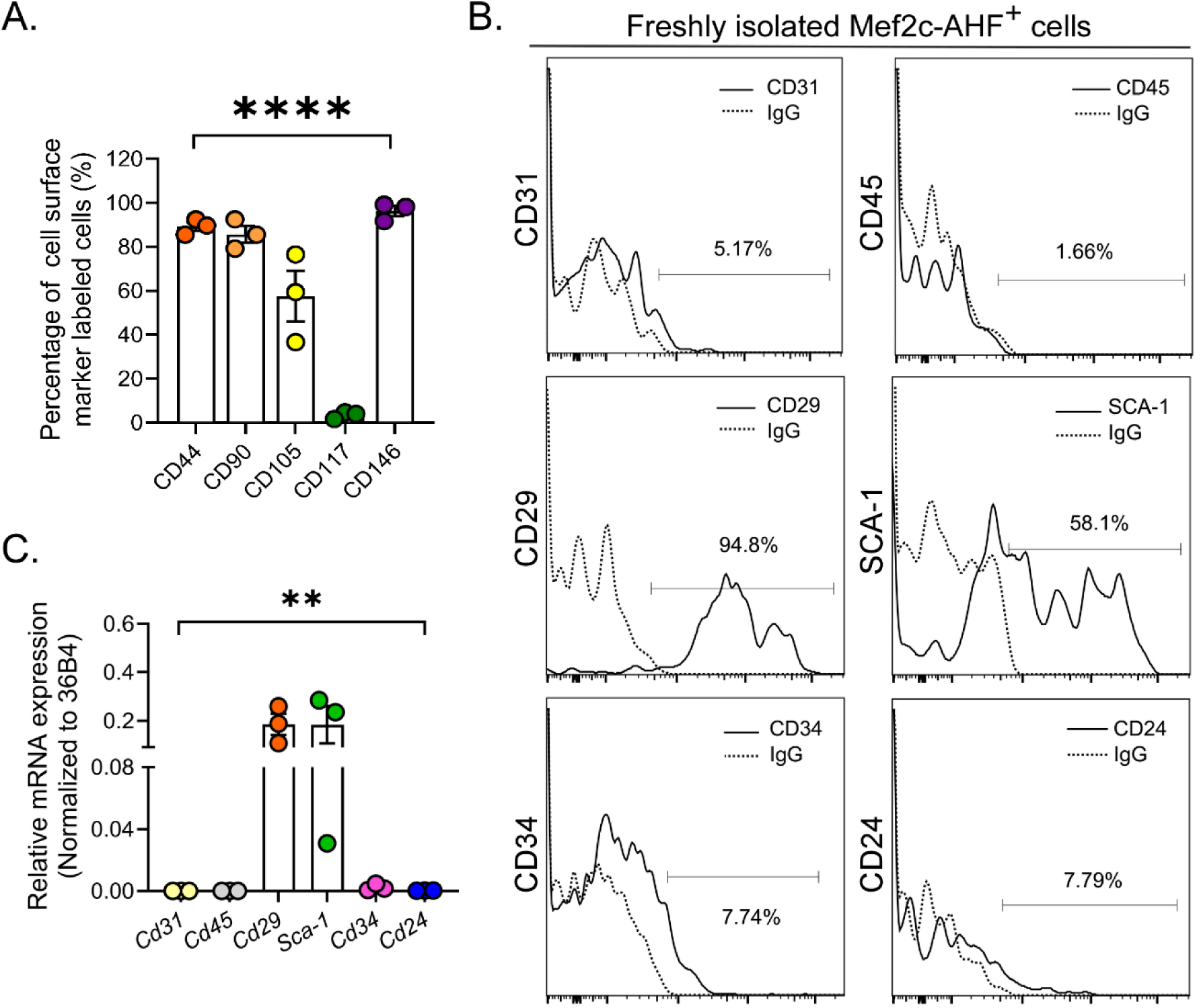
Cell surface marker analyses of freshly isolated Mef2c-AHF^+^ cells. (A) Quantification of the percentage of cells positive for CD44, CD90, CD105, CD117, and CD146 in the Mef2c-AHF^+^ cell population compared to the relevant isotype control. Data are presented as mean ± SEM. ****, p < 0.0001. n = 3. (B) Representative histogram plots showing the expression of CD31, CD45, CD29, SCA-1, CD34, and CD24 (solid line) with the relevant isotype control (dashed line, IgG) in freshly isolated Mef2c-AHF^+^ cells from the scBAT of postnatal 3-week-old *Mef2c-AHF-Cre;ROSA^mTmG^* mice. n = 2. (C) Relative mRNA levels of *Cd31*, *Cd45*, *Cd29*, *Sca-1*, *Cd34*, and *Cd24* in Mef2c-AHF^+^ cells isolated from the scBAT of postnatal 3- to 5-week-old *Mef2c-AHF-Cre;ROSA^mTmG^* mice. Data are presented as mean ± SEM. **, p < 0.01. n = 3.

**Figure S5.**
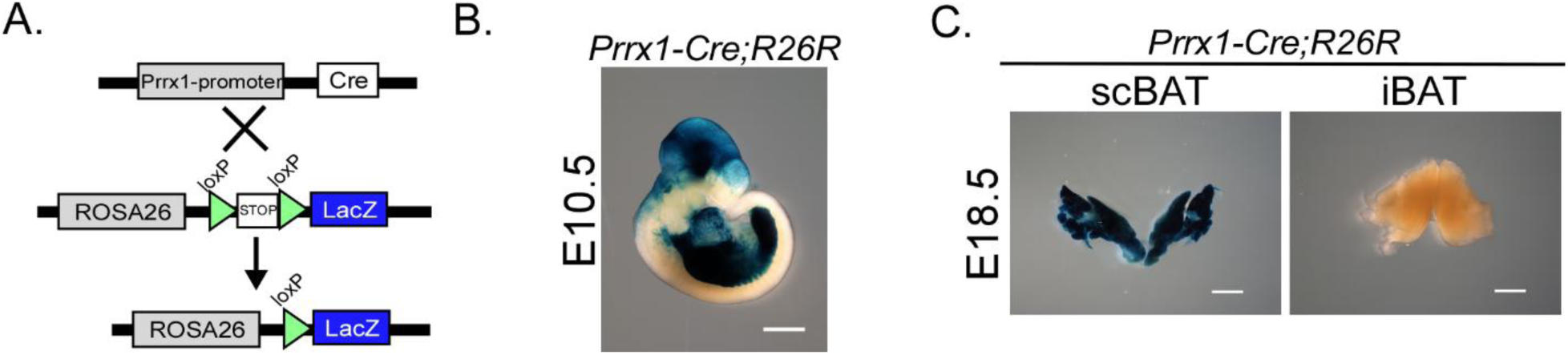
A Prrx1-Cre driver marks scBAT in mouse embryos. (A) A schematic diagram of the lineage tracing strategy using *Prrx1-Cre* mice crossed with *R26R* mice to identify Prrx1- marked BAT. (B) Representative image of whole mount X-gal-stained embryos from E10.5 *Prrx1- Cre;R26R* embryos. Scale bar = 2000 μm. n = 3. (C) Representative images of X-gal-stained scBAT and iBAT from E18.5 *Prrx1-Cre;R26R* embryos. Scale bar = 2000 μm. n = 3.

**Figure S6.**
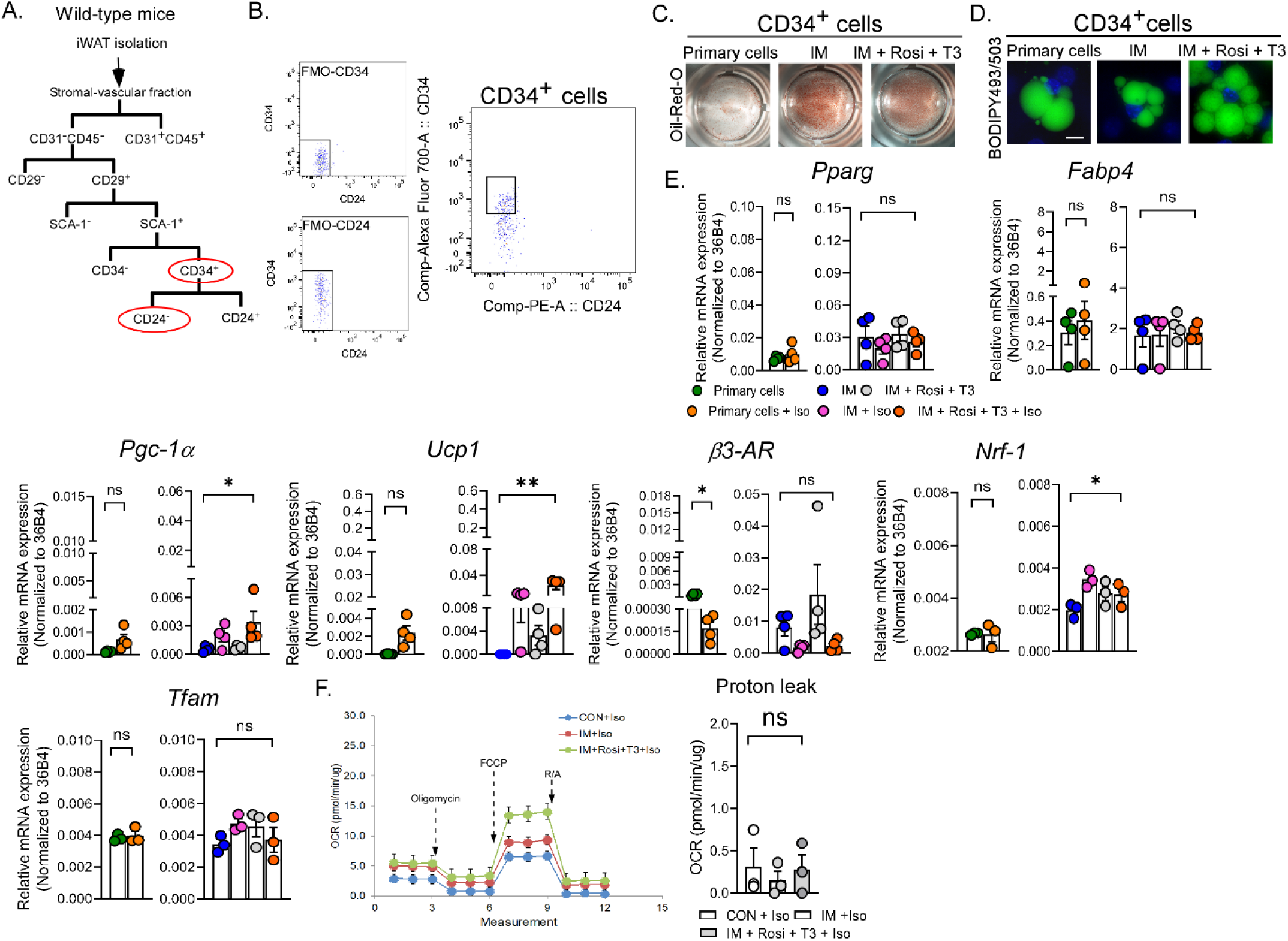
CD34^+^ cells isolated from the iWAT of wild-type mice can differentiate into white adipocytes. (A) A schematic diagram of the FACS strategy for isolating CD34^+^ cells from the iWAT of postnatal wild-type mice. (B) Representative density plots showing FACS staining profiles and gating (black boxes) for the isolation of CD34^+^ cells. n = 50. FMO: Fluorescence Minus One. (C) Representative images of Oil-Red-O-stained CD34^+^ cells that differentiated spontaneously or were induced by IM or by a combination of IM, Rosi, and T3. n = 4. (D) Representative fluorescent images of BODIPY493/503-stained CD34^+^ cells that differentiated spontaneously or were induced by IM or by a combination of IM, Rosi, and T3. Scale bar: 10 μm. n = 3-5. (E) Relative mRNA levels of genes involved in beige adipogenesis and mitochondrial biogenesis that differentiated spontaneously or were induced by IM or by a combination of IM, Rosi, and T3. Iso was added at the end of culture period. Data are presented as mean ± SEM. **, p < 0.01; *, p < 0.05; ns = nonsignificant. n = 3-4. (F) Representative Seahorse assay traces and a graph of calculated protein leak in CD34^+^ cells that differentiated spontaneously (CON) or were induced by IM or by a combination of IM, Rosi, and T3. Iso was added to stimulate thermogenesis at the end of the culture period. OCR values were normalized to the total protein amount. Data are presented as mean ± SEM. ns = nonsignificant. n = 3.

**Figure S7.**
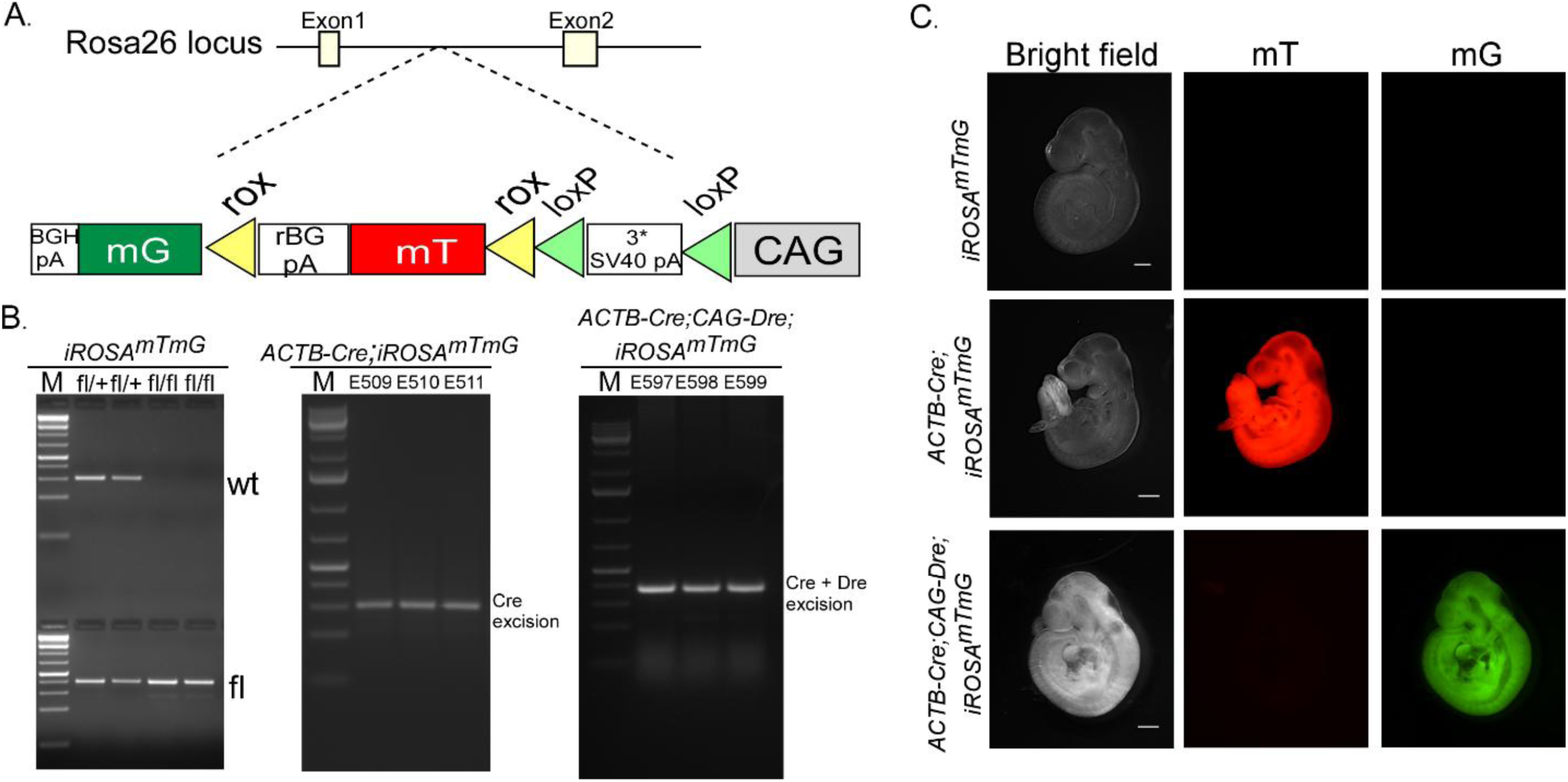
The generation of intersectional ROSA^mTmG^ mice (*iROSA^mTmG^*). (A) A schematic diagram of the gene targeting vector design. The *iROSA^mTmG^*reporter mouse was generated by knocking a targeting vector consisting of a ubiquitous CAG promoter, loxp-flanked Stop codons, a rox-flanked membrane-targeted tdTomato (mT) gene, and a membrane-targeted GFP (mG) gene into the *ROSA26* locus in the antisense orientation. (B) Representative images of PCR genotyping results. These images include PCR products indicating homozygous *iROSA^mTmG^*mice, generated by breeding *iROSA^mTmG^* heterozygous male and female mice (left). The PCR products indicated that ACTB-Cre-mediated excision of the stop codons in the *ACTB-Cre;iROSA^mTmG^* pups (middle). These pups were generated by breeding *ACTB-Cre* male mice with homozygous *iROSA^mTmG^* female mice. The PCR products indicate that the CAG-Dre-mediated excision of the mT in *ACTB-Cre;CAG-Dre;iROSA^mTmG^* pups (right). These pups were generated by breeding *ACTB-Cre;CAG-Dre* double heterozygous male mice with the homozygous *iROSA^mTmG^* female mice. fl: flox. (C) Bright field and fluorescent images of E9.5 embryos of *iROSA^mTmG^*, *ACTB-Cre;iROSA^mTmG^*, and *ACTB-Cre;CAG-Dre;iROSA^mTmG^*. In this reporter mouse, neither mT nor mG is expressed in the absence of Cre. mT is expressed in the presence of the Cre, while mG is expressed in the presence of both Cre and Dre. Scale bar = 2500. μm. n = 3-4.

## Methods

### Materials Availability

Most of the mouse lines used in this study were previously published by other laboratories. The new mouse and cell lines generated in this study can be provided upon request, following the completion of a Materials Transfer Agreement.

### Data Availability

The RNA sequencing datasets that were reported in this paper have been deposited in the NIH Gene Expression Omnibus (GEO).

## Experimental Details

### Mouse models

Wild-type mice (C57BL/6J) were purchased from the Center for Comparative Medicine (CCM) production colony at BCM. Genetically modified mice include *ROSA^mTmG^* mice^48^ (stock #007676), *R26R* mice^49^ (stock #003474), *Myf5^Cre^*mice^50^ (stock #007893), *Pax7^Cre^* mice^51^ (stock #010530), *Pax3^Cre^*mice^52^ (stock #005549), *Prrx1-Cre* mice^41^ (stock #005584), *ACTB-Cre* mice^53^ (stock#019099), and *Ucp1-Cre* mice^54^ (stock #024670), which were purchased from the Jackson Laboratory. *Mef2c-AHF-Cre* mice^34^ (MMRRC, 030262-UNC) and *CAG-Dre* mice^55^ (MMRRC, 032246-UCD) were rederived by the BCM Genetically Engineered Mouse Core with frozen sperm purchased from MMRRC. The *Mef2c-AHF-DreERT2* and *ROSA^nKmB^* mice^35^ were generously provided by Dr. Benoit Bruneau (Gladstone Institutes, San Francisco, CA, USA) through Dr. Joshua Wythe (University of Virginia School of Medicine, VA, USA) and were rederived by the BCM Genetically Engineered Mouse Core. The intersectional ROSA^mTmG^ (*iROSA^mTmG^*) mice were generated through ES cell-mediated gene targeting, using a fee-based commercial service (Cyagen US Inc., CA, USA). All animal experiments were approved by the Institutional Animal Care and Use Committee (IACUC) at BCM and conducted in accordance with its guidelines. All the mice used in this study were housed on a standard 14-hour light/10-hour dark cycle at ambient temperature, with unrestricted access to standard rodent chow and water. Genetically modified mice were genotyped using primers listed in Table S1.

## METHOD DETAILS

### Tissue isolation, processing for cryosection, and imaging

Adipose tissues, including scBAT, iBAT, iWAT, and eWAT, were collected from mouse embryos or from pups (3-5-week-old). Transcardiac perfusion with 4% paraformaldehyde was performed prior to the collection of adipose tissue from the pups. The collected adipose tissues were subsequently fixed in 4% paraformaldehyde at 4°C overnight, cryopreserved with sucrose, and embedded in optimal cutting temperature compound (OCT). The embedded adipose tissue was sectioned at 10-15 µm thickness using a Leica cryostat and imaged using a Nikon Eclipse 80i stereomicroscope. E9.5 and E10.5 embryos and E12.5 hearts were dissected and imaged using a Nikon SMZ18 stereomicroscope.

### Whole mount X-gal staining

Whole mount X-gal staining was performed according to a previously published procedure with slight modifications^56^. Briefly, E10.5 embryos, E12.5 hearts, or E18.5 BAT depots were collected and fixed in a fixative solution containing 2% paraformaldehyde, 0.2% glutaraldehyde, 5 mM EGTA, and 2 mM MgCl2 at 4°C for a duration ranging from 2 hours to overnight. Embryos, hearts, or BAT depots were then rinsed three times in Phosphate Buffered Saline (PBS) and stained for β-galactosidase overnight in the dark at room temperature in staining solution, consisting of 5 mM potassium ferrocyanide, 5 mM potassium ferricyanide, 2 mM MgCl2, 1 mg/ml 5-bromo-4-chloro- 3-indolyl-β-D-galactopyranoside (X-gal), 0.02% IGEPAL, 0.01% sodium deoxycholate, and 20 mM Tris, pH 7.4. Following staining, embryos, hearts, or BAT depots were rinsed twice in PBS and post-fixed in 4% paraformaldehyde overnight at 4°C. X-gal-stained E10.5 embryos, E12.5 hearts, and E18.5 BAT depots were cleared in a series of glycerol/PBS solutions (20%, 50%, and 80%) and imaged using a Nikon SMZ1500 stereomicroscope.

### Tamoxifen (Tam) treatment

To induce Dre recombinase activity, mice were intraperitoneally injected with Tam (60-80 mg/kg body weight) dissolved in corn oil.

### Adipocyte quantification

To quantify the Mef2c-AHF positive (mG-labeled) adipocytes, scBAT was collected and embedded in OCT. For adipocyte counting, 10 µm transverse frozen sections were imaged using a Nikon 80i stereomicroscope and processed with NIS Elements Software (AR 3.2). The software was utilized to isolate tdTomato (mT) and GFP (mG) channels. After subtracting the background, the adipocyte cell boundary was determined by visually identifying the intensity of fluorescence for each cell. Mef2c-AHF positive adipocytes were manually counted using the “Counts” feature found under Annotations and Measurements in NIS Elements. The percentage of GFP expression was quantified as the number of GFP positive adipocytes over the total number of adipocytes.

### Isolation and immortalization of stromal-vascular fraction (SVF) cells from adipose tissue

The SVF was isolated from scBAT or iWAT using a previously published method with some minor modifications^12^. Briefly, the isolated adipose tissues were rinsed with PBS, minced into small pieces with a blade, and digested in a freshly made digestion medium. The digestion medium consisted of 0.5 mg/ml collagenase A, 1.2 u/ml dispase II, and 20 u/ml DNase I in Seahorse XF based medium (Agilent) supplemented with 25 mM glucose, 1 mM sodium pyruvate, and 2 mM L-glutamine. Antibiotics, including 1X Pen/Strep, 1X gentamicin, and 1X amphotericin B were added to the digestion medium prior to use. The tube containing the minced adipose tissues was placed in a shaker and incubated at 37°C for 40-60 minutes, shaking at 200 rpm. After digestion, 10 ml of fresh medium (high-glucose DMEM, 10% EqualFetal bovine serum, Pen/Strep, and gentamicin) was added to the tube to neutralize the digestion mixture. The mixture was then filtered through a cell strainer (40 µm diameter) to remove any undigested debris, and centrifuged at 300 g for 10 minutes to pellet the SVF. The pelleted SVF cells were incubated with red blood cell lysis buffer (Miltenyi Biotec.) for 5 minutes and centrifuged at 1500 rpm for another 5 minutes to remove red blood cells. Some SVF cells isolated from scBAT were immortalized with SV40 large T antigen following a previously published procedure^57^.

### Flow cytometry

For flow cytometric analysis, the pelleted SVF cells from the scBAT of *Mef2c-AHF-Cre; ROSA^mTmG^* pups (5-7-day-old or 3-week-old) were resuspended in 1 ml of Opti-MEM. The number of SVF cells was counted using a Cellometer Mini (Nexcelom) or a hemocytometer. After counting, the SVF cells were washed once with Opti-MEM, and at least 1x10^5^ live SVF cells were used for antibody staining at 4°C for 30 minutes. The stained SVF cells were washed once and resuspended in PBS for flow cytometric analysis using an LSR II Flow Cytometer (BD Biosciences) at the Texas Children’s Hospital Flow Cytometry Core Laboratory. Wild-type cells, unstained SVF cells, and isotype controls were used as gating controls. Flow cytometry plots were generated using FlowJo software (V10). Antibodies used for the flow cytometric analysis included CD45 (Biolegend, #103111, 1:400), CD31 (Biolegend, #102409, 1:400), CD29 (Biolegend, #102215, 1:400), SCA-1 (Biolegend, #108111, 1:400), CD24 (Biolegend, #138505, 1;200), CD34 (Biolegend, #128611, 1:100), Rat IgG2a, k Isotype (Biolegend, #400511, 1:400), Rat IgG2b, k Isotype (Biolegend, #400611, 1:400), Rat IgG2c, k Isotype (Biolegend, #400713, 1:200), Armenian Hamster IgG Isotype (Biolegend, #400911, 1:100), CD44 (Biolegend, #103011, 1:400), CD90.2 (Biolegend, #105311, 1:400), CD117 (Biolegend, #105811, 1:100), CD105 (Biolegend, #120413, 1:400), CD146 (Biolegend, #134711, 1:400).

### Fluorescence-activated cell sorting (FACS)

For sorting immortalized Mef2c-AHF^+^ cells, immortalized SVF cells from scBAT of the *Mef2c- AHF-Cre; ROSA^mTmG^*pups (5 days old) were trypsinized in 0.25% Trypsin-EDTA (Gibco) and resuspended in a sorting buffer. This buffer consisted of 1X Hanks’ Balanced Salt Solution (HBSS), 3% BSA, 25 mM HEPES, and antibiotics (Pen/Strep, gentamicin, amphotericin B) buffered to pH 7.4. Mef2c-AHF-marked cells (GFP positive) were collected for use in the subsequent experiments. For sorting lineage-traced primary cells, the pelleted SVF cells from scBAT of the *Mef2c-AH-FCre;ROSA^mTmG^*or *Prrx1Cre;ROSA^mTmG^* pups (2-5-week-old) were washed and resuspended in the sorting buffer. Mef2c-AHF- or Prrx1-marked cells (GFP positive) were collected for use in subsequent experiments. For sorting wild-type primary cells, the pelleted SVF cells from scBAT or iWAT of the C57BL/6J pups (2-5-week-old) were washed once with the sorting buffer and stained with antibodies at 4°C for 30 minutes. After staining, the SVF cells were washed once and resuspended in the sorting buffer. CD34^-^ and CD34^+^ cells were collected for use in subsequent experiments. On average, adipose tissues from 5-6 pups were collected for each sorting experiment. FACS gating was established using wild-type cells and fluorescence minus one (FMO) controls. All the sorted cells were collected in a collection buffer consisting of Seahorse XF based medium supplemented with 25 mM glucose, 1 mM sodium pyruvate, 2 mM L-glutamine, antibiotics, and 50% EqualFetal bovine serum. FACS was conducted using a BD Biosciences FACSAria cell sorter at the Texas Children’s Hospital Flow Cytometry Core Laboratory. Flow cytometry plots were generated using FlowJo software (V10). The antibodies used for FACS included CD45 (Biolegend, #103107, 1:1000), SCA-1 (Biolegend, #108119, 1:200), CD24 (Biolegend, 138503, 1:200), CD31 (Biolegend, #102417, 1:200), CD34 (BD BioSciences, #560518, 1:50), and CD29 (Biolegend, #102215, 1:200).

### BMP7 conditioned medium production

To produce BMP7 conditioned medium, a mammalian Bmp7 expression plasmid, PRK5::Bmp7 (10 μg), was transiently transfected into HEK293T cells (10 cm plate) using Lipofectamine 3000 according to the manufacturer’s instructions (Thermo Fisher Scientific). The conditioned medium was collected 48 hours after transfection, filtered through a 22 µm filter, and stored in a -80°C freezer.

### Adipocyte progenitor cell culture and differentiation

After sorting, adipocyte progenitor cells (APCs) were seeded in the primary cell basal medium in a 24-well plate. This medium consisted of 60% low glucose DMEM, 40% MCDB201, 1% Insulin-Transferrin-Selenium (ITS), 1% linoleic acid-BSA, 10 μM 2-phospho-L-ascorbic acid, and antibiotics (Pen/Strep, gentamicin, amphotericin B), buffered to pH 7.4. Growth factors, including 10 ng/ml bFGF, 10 ng/ml EGF, and 1x10^3^ LIF, along with 10% EqualFetal serum, were added to the basal medium prior to cell culture. After the first passage, the sorted APCs continued to be cultured in the basal medium supplemented with 4% EqualFetal serum and growth factors. APCs were expanded once or twice to obtain enough cells for the subsequent treatments. For IM treatment, APCs isolated from scBAT were cultured in basal medium supplemented with 2% EqualFetal serum, antibiotics, 0.5 mM 3-isobutyl-1-methylxanthine (IBMX), 1µM Dexamethasone, 5 µg/ml Insulin, and 1 nM 3,3′,5-Triiodo-L-thyronine (T3) for 48 hours and maintained in medium containing Insulin and T3 until fully differentiated (approximately 6-8 days after induction). For BMP7 and rosiglitazone treatment (BR), 1 µM Rosiglitazone and 25% (v/v) of BMP7 supernatants were added to the basal medium supplemented with 2% EqualFetal serum and antibiotics. Thermogenesis was induced by incubating with 5 µM Isoproterenol (Iso) for 6 hours. APCs isolated from iWAT were induced with the same induction cocktail without T3. To induce beiging, 1 µM Rosiglitazone and T3 were included in IM. Immortalized Mef2c-AHF^+^ cells were cultured in culture medium containing high-glucose DMEM, 10% EqualFetal serum, Pen/Strep, and gentamicin supplemented with 50-100 µg/ml G418.

### RNA isolation and RT-qPCR

Total RNA was extracted using a RNeasy Micro Kit (Qiagen) or an Aurum Total RNA Fatty and Fibrous Tissue Kit (Bio-Rad), following the manufacturer’s instructions. 250 ng of total RNA was used to synthesize first-strand cDNA using the SuperScript III First-Strand Synthesis System (Thermo Fisher Scientific). Quantitative PCR reactions were performed using Power SYBR Green PCR Master Mix (Thermo Fisher Scientific) in a CFX96 Touch Real-Time PCR Detection System or CFX Opus 96 system (Bio-Rad). The ΔCt method (2^-ΔCt^) was used to calculate the relative mRNA expression level of each gene. Specific gene expression was normalized to either *β-actin* or *36B4*. Primers used for RT-qPCR are listed in Table S2.

### Oil-Red-O staining

The sorted APCs were differentiated in a 48-well plate. After differentiation was complete, the adipocytes were rinsed in PBS and fixed in 4% paraformaldehyde for 30 minutes. After fixation, the adipocytes were washed once in ddH2O, followed by a brief rinse in 60% isopropanol, and stained with Oil-Red-O solution at room temperature for 2 hours. The stained adipocytes were rinsed twice in 60% isopropanol and once or twice in ddH2O to remove excess Oil-Red-O dye. The Oil-Red-O-stained adipocytes were then imaged using a Nikon SMZ1500 stereomicroscope.

### LipidSpot 610 staining

The sorted APCs were cultured on coverslips coated with collagen coating solution (Sigma) in a 48-well plate. After differentiation was complete, the differentiated adipocytes were rinsed once in PBS and fixed in 4% paraformaldehyde for 1 hour. The fixed adipocytes were permeabilized in 0.2% Triton X-100/PBS for 15 minutes, washed three times in 0.02%Triton X-100/PBS, and stained with LipidSpot 610-diluted in 0.02% Triton X-100/PBS (1:1000 dilution) for 1 hour. The stained adipocytes were further washed three times in 0.02% Triton X-100/PBS and counterstained with DAPI for 10 minutes. After staining was complete, the coverslips were removed from the plate and placed invertedly onto a microscope slide coated with a drop of antifade mounting solution (Vector laboratories). The LipidSpot 610-stained adipocytes were imaged using a Nikon Eclipse 80i stereomicroscope.

### BODIPY 493/503 staining

The sorted APCs were cultured on coverslips coated with collagen coating solution (Sigma) in a 48-well plate. After differentiation was complete, the adipocytes were rinsed once in PBS and fixed in 4% paraformaldehyde for 30 minutes. The fixed adipocytes were washed once more with PBS and stained with BODIPY 493/503 diluted in PBS (1:1000 dilution) for 30 minutes in the dark. Following staining, the adipocytes were washed 3 times in PBS and counterstained with DAPI for 10 minutes. After staining was complete, the coverslips were removed from the plate and placed inverted onto a microscope slide coated with a drop of antifade mounting solution. The BODIPY 493/503-stained adipocytes were imaged using a Nikon Eclipse 80i stereomicroscope.

### Protein isolation and western blotting

Protein isolation was performed as previously described^12^. Protein concentrations were measured using a Pierce BCA Protein Assay Kit (Thermo Fisher Scientific). For western blotting, 10-30 μg of protein lysates were subjected to SDS-PAGE and transferred onto Immun-Blot PVDF membranes (Bio-Rad) using an eBlot L1 Fast Wet Transfer System (GenScript), following the manufacturer’s instructions. The transferred membranes were blocked in 5% milk/TBST buffer (50 mM Tris pH 7.5, and 150 mM NaCl) for 1 hour or overnight at 4°C. The membranes were then incubated with primary antibodies in 3-5% BSA/TBST or 5% milk/TBST for 2 hours at room temperature or overnight at 4°C and washed 4 times in TBST. After washing, the membranes were incubated with secondary antibodies in 2.5% milk/TBST for 1 hour at room temperature and washed 4 times in TBST. The stained membranes were developed using Pierce ECL Plus Western Blotting Substrate according to the manufacturer’s instructions (Thermo Fisher Scientific). The antibodies used for western blotting included PPARγ (Cell Signaling, #2430S, 1:1000 or Santa Cruz, #sc-7273, 1:500), FABP4 (Abcam, 13979, 1:2000), UCP1 (Abcam, #10983, 1:2000), BMP7 (Cell Signaling, #4693S, 1:1000), BMPR1A (Proteintech, #12701-1-AP, 1:1000), BMPR2 (Cell Signaling, #6979S, 1:1000), SMAD1 (Cell Signaling, #9743S, 1:1000) and β-ACTIN (Cell Signaling, #4976S, 1:2000), and HRP (Jackson ImmunoResearch, Code#715-035-150, 1:3000 or Code#715-035-152, 1:3000).

### Mitochondrial respiration measurement

The freshly sorted primary cells were seeded onto a XF24 cell culture microplate (Agilent) at 1x10^4^ cells per well. The cells were cultured with or without adipogenic inducers for 8 days. Some of the cells were incubated with Isoproterenol (Iso) (5 µM) for 4 hours at the end of the culture period. To measure mitochondrial respiration, cells were rinsed (1 ml per well) and incubated (500 µl per well) in Seahorse XF basal medium (pH 7.4) containing 10 mM glucose, 2 mM L-glutamine, and 1 mM sodium pyruvate for 1 hour at 37°C in a non-CO2 incubator. After incubation, the microplate was loaded onto the Seahorse XFe24 Analyzer (Agilent). The oxygen consumption rate (OCR) was measured three times at three-minute intervals at baseline, 3 times after the injection of Oligomycin (2 µM), 3 times after the injection of FCCP (5 µM), and finally 3 times after the injection of rotenone and antimycin (R/A, 0.5-2 μM each). The OCR of each sample was normalized to the total protein content of the sample and analyzed using the Seahorse XF Cell Mito Stress Test Report Generator. Data from 3 individual assays were combined using the multi-file Seahorse XF Cell Mito Stress Test Report Generator (Agilent).

### RNA-seq analysis

RNA-seq was performed using the immortalized Mef2c-AHF^+^ cells that were either not differentiated (CON) or differentiated with IM or BR for 2 days. Total RNA from each sample was extracted using an Aurum Total RNA Fatty and Fibrous Tissue Kit (Bio-Rad). Samples from two independent experiments were sent to the BCM Children’s Nutrition Research Center Laboratory for Translational Genomics for quality checks, library construction, and sequencing. The raw sequencing data containing pair-ended reads were mapped to the mouse genome (USCS mm10) using software STAR^58^ with NCBI RefSeq genes as the reference. STAR generated read counts for each gene were used as the measurement for gene expression. DESeq2^59^ was used to analyze the gene-based read counts to detect differentially expressed genes between the groups of interest. The hierarchical clustering tree was generated by inputting the normalized gene expression levels (Reads Per Million, RPM) of the CON, IM, or BR-treated Mef2c-AHF^+^ cells into MeV (version 4.9.0), followed by Phylip (version 3.698) DrawTree software. The final hierarchical clustering tree image was created in Adobe Illustrator. Venn diagrams (Venny 2.0) were generated by comparing genes that were up-regulated or down-regulated (FDR value < 0.01 and fold change > 2) in IM or BR-treated Mef2c-AHF^+^ cells compared to CON cells. GO term analysis of these up-regulated or down-regulated genes was performed using WebGestaltR (webgestalt.org). The top 5 GO terms were included in Suppl Fig.3. The comparison of the expression of genes involved in adaptive thermogenesis (Gene Ontology term (GO): 1990845) in IM or BR-treated Mef2c-AHF^+^ cells, as well as the generation of the heatmap, was performed using RStudio.

### Quantification and Statistical Analysis

The student’s t-test was used to evaluate statistical significance between 2 groups. One-way ANOVA was used to evaluate statistical significance between 3 or more groups. P<0.05 was considered statistically significant. Statistical analyses were performed using GraphPad Prism 10 software.

---

**Table S1:**
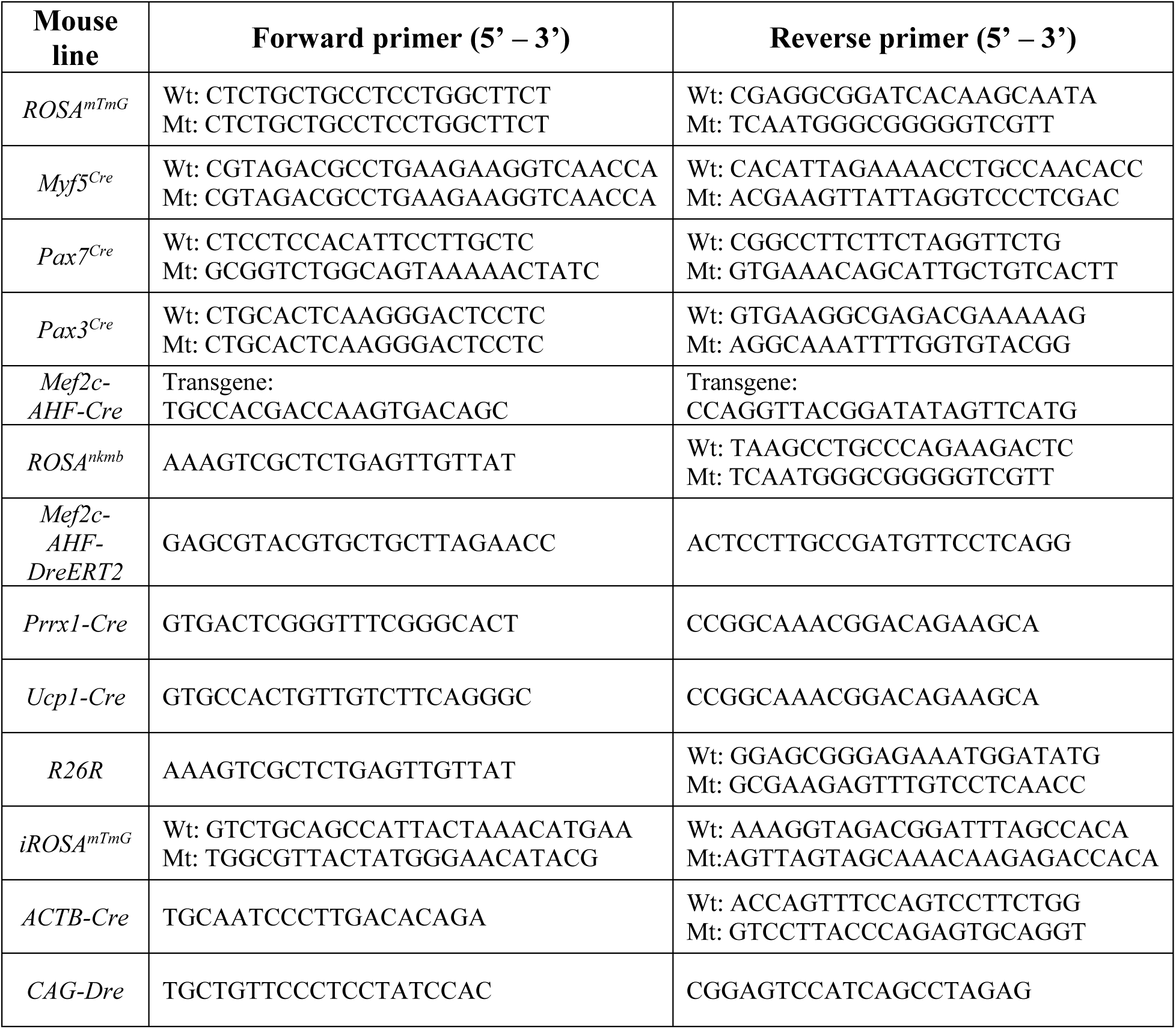
Primer sequences for mouse genotyping.

**Table S2:**
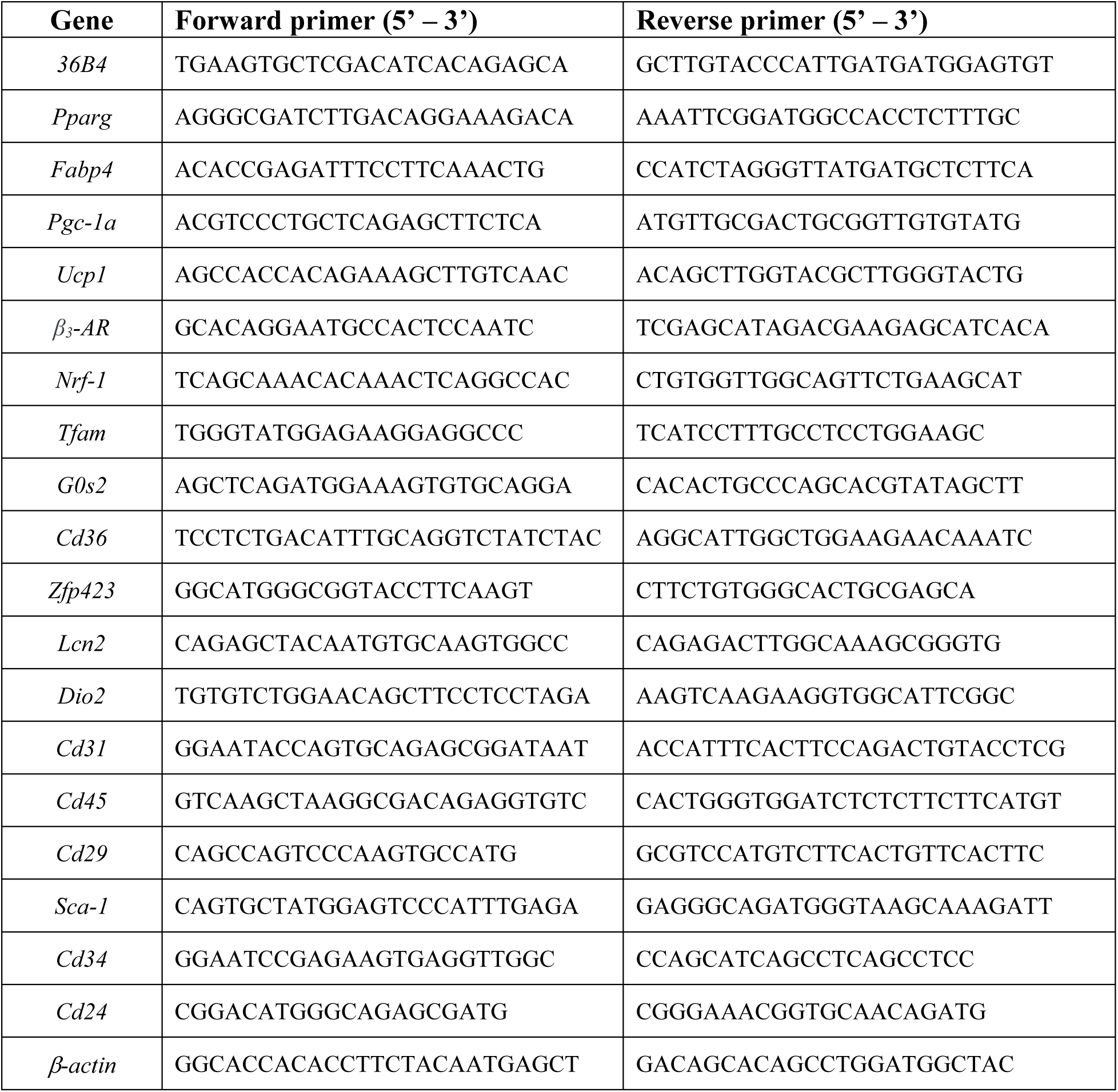
RT-qPCR primer sequence used in this paper.

